# Engineered flavonoid disrupts mitochondrial AIF/CHCHD4 complex for targeted cancer therapy

**DOI:** 10.1101/2025.03.13.642976

**Authors:** Hong Toan Lai, Daniela Dias - Pedroso, Maria Eugénia Marques Da Costa, Romain Fernandes, Tran Ngoc Anh Nguyen, Olivia Bawa, Pierre Khneisser, Svetlana Dokudovskaya, Guido Kroemer, Antonin Marchais, Nathalie Gaspar, Birgit Geoerger, Oumar Diane, Liuba Mazzanti, Tâp Ha Duong, Guy Lewin, Laurent Ferrié, Bruno Figadère, Catherine Brenner

## Abstract

Oncogenic metabolism depends on multifaceted mechanisms, including bidirectional inter-organelle communication between mitochondria and the nucleus, facilitating cellular adaptation at the transcriptomic, proteomic, and metabolomic levels. The mitochondrial protein complex composed of apoptosis-inducing factor (AIF) and coiled-coil-helix-coiled-coil-helix domain-containing protein 4 (CHCHD4) is essential for this mitochondrio-nuclear communication. The AIF/CHCHD4 complex mediates the mitochondrial import of cysteine-enriched nuclear gene-encoded proteins, thereby adapting the mitochondrial proteome to cellular energy demands. We report the discovery of M30-E05, an engineered flavonoid that binds to the NADH pocket of AIF, preventing its dimerization and disrupting the AIF/CHCHD4 complex. Molecular docking and gel electrophoresis analysis of mitochondrial AIF/CHCHD4 substrates expression confirm this mechanism. In cancer cells, M30-E05 reduces the expression of nuclear gene-encoded mitochondrial proteins such as AIF, CHCHD4, COX17, and MICU1. In addition, M30-E05 fragments the mitochondrial network and impairs mitochondrial respiration, causing profound alterations, particularly in lipid and aminoacid metabolism, as revealed by kinetic measurements of oxygen consumption and mass spectrometric metabolomics. Importantly, M30-E05 significantly reduces the viability of a human adult and pediatric osteosarcoma cancer cell panel, including those from patient-derived xenografts (PDX) of osteosarcomas, and induces apoptosis. When orally administered for two weeks to immunodeficient NSG mice, M30-E05 inhibited tumor growth in a subcutaneous PDX xenograft model without apparent toxicity. We anticipate that M30-E05, as a first-in-class metabolic inhibitor, could serve as the lead compound for a new class of targeted antineoplastic agents.

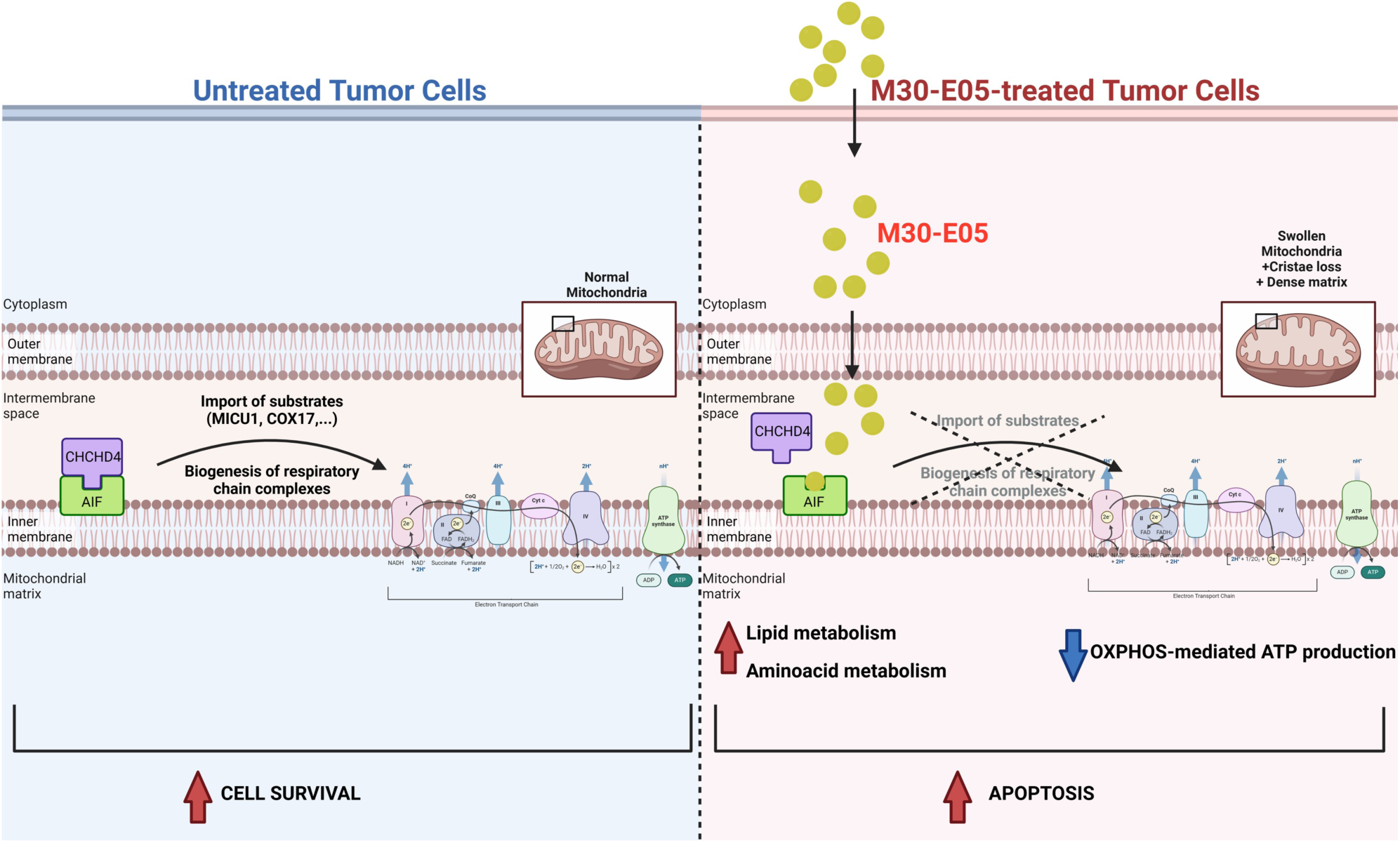

## Introduction

The first theory of oncogenic metabolic reprogramming proposed by Otto Warburg^1^ describes a shift in ATP production from mitochondrial oxidative phosphorylation (OXPHOS) to glycolysis, which may contribute to the higher proliferative rate of cancer cells. Today, the concept of metabolic reprogramming has expanded beyond energy metabolism and includes explanations for the formidable capacity of cancer cells to engage in anabolic reactions. Advances in our understanding of tumor biology and its interaction with the microenvironment have led to the recognition of dysregulated cellular metabolism as a new hallmark of cancer^2,3,4^. Indeed, in cancer patients, hypermetabolism, which can be detected by PET scan imaging, is typically associated with poor survival rates and can predict cachexia and relapse. Cancer cells exploit their intrinsic metabolic flexibility, enabling them to switch between different energy substrates and to equilibrate anabolism and catabolism to survive within the tumor microenvironment. Building on these observations, we hypothesized that small molecules that subvert this metabolic flexibility could be used to inhibit tumor progression^5^.

Oncogenic metabolism depends on bidirectional inter-organelle communication between mitochondria and the nucleus, which allows cellular adaptation at the transcriptomic, metabolomic, and proteomic levels, ultimately promoting cancer progression^6^ ^7^. Most of ∼1,500 mitochondrial proteins are encoded by the nuclear genome and imported into the mitochondria^8^. To meet cancer cell-specific needs, several specialized protein import machineries have evolved, adapting the mitochondrial proteome accordingly. Within these import pathways, the disulfide relay machinery relies on the formation of a complex between apoptosis-inducing factor (AIF, encoded by *AIFM1* gene) and coiled-coil-helix domain-containing protein 4 (CHCHD4) complex. This complex regulates the redox-sensitive import of small, cysteine-containing proteins, critical for cell survival and stress adaptation^9^ ^6^. Monomeric, dimeric AIF and CHCHD4 contribute to mitochondrial bioenergetics, redox regulation, lipid and calcium homeostasis, and to the maintenance of mitochondrial^6^. Genetic alterations affecting AIF influence mitochondrial biology and cell fate through the biogenesis of the respiratory chain and the import of key nuclear-encoded mitochondrial proteins, such as MICU1, C9orf72, NDUFS5 and TRIAP1^10^ ^11^ ^12^ ^13^. Loss-of-function mutations in *AIFM1* are linked to a range of X-linked mitochondrial diseases^14^. In contrast, the potential oncogenic role of unmutated *AIFM1* has been demonstrated in a mouse model of KRAS-mutant lung cancer^15^. Similarly, CHCHD4 favors tumor proliferation and epithelial-to-mesenchymal transition (EMT)-related phenotypes through respiratory chain-mediated metabolism^16^.

To test whether disrupting the AIF/CHCHD4 protein complex could reduce metabolic flexibility and promote cancer cell death, we designed an original highly specific, high-throughput bioassay to identify inhibitors disrupting this complex^17^. Using this assay, we screened several libraries, including the Prestwick Library of FDA-approved drugs, a subset of the French National Chemical Library, and a BioCIS collection of new molecules with diverse chemical structures. Notably, this BioCIS collection included derivatives of flavonoids, chemically modified natural compounds, that show promise for therapeutic applications with minimal toxicity. As a result, the screening identified several potential AIF ligands (i.e., “hits”) that were ranked based on their ability to disrupt the AIF/CHCHD4 complex. These hits were further evaluated for their pharmacological activity through *in silico* analysis, *in cellulo* models including both parental and drug-resistant cancer cell lines, secondary cultures of osteosarcoma patient-derived xenografts (PDX) as well as *in vivo* PDX models.

Our findings show that M30-E05 targets oncogenic metabolism and induces apoptosis by disrupting the AIF/CHCHD4 complex in pediatric and adult cancer cell line panel, significantly reducing osteosarcoma tumor growth *in vivo*. Given that metabolic reprogramming is a universal feature of cancer, M30-E05 represents a promising lead for the development of a novel class of targeted antineoplastic agents.

## Materials and Methods

### Chemicals

M30-E05 has been synthesized in BioCIS laboratory (details provided in Supplementary Notes). The compound was solubilized in dimethyl sulfoxide (DMSO) at 25 mM or 50mM stock solutions, and stored at –20 °C. For *in vitro* experiments, M30-E05 was washed with a 1:1 Chloroform: Methanol solution, air-dried overnight using by Speedvac, before solubilization. For *in vivo* studies, the compound was solubilized in 0.5% methylcellulose (Merck, M7027), freshly before use.

### Screening on inhibition of interaction between HIS-AIF_103-613_ and GST-CHCHD4

HIS-AIF_103-613_ and GST-CHCHD4-Gst were produced in BL21+DE3(RIPL) bacteria during a 3h-induction with 0.5 mM IPTG at 37°C, as previously described^18^. Assays were performed in white 384-well microplates (Greiner, 781075) with a total volume of 25 µL per well. Proteins, peptide N27 (positive control)^18^ and beads were diluted in 0.1 % Bovine Serum Albumin (BSA)/Phosphate-buffered Saline (PBS), on ice.

A total of 1.280 off-patent small molecule library provided by Prestwick Chemicals; 2.901 compounds provided by the French National Library and 99 compounds provided by BioCIS library were screened at 10 µM in 0.1% DMSO in monoplicate. Firstly, 5 μL of 10 µM compound or 0.1 % DMSO / 3 µM N27 or 0.1 % DMSO / 0.1 % BSA PBS (negative control) were added. Secondly, 5 µL of His-AIF_103-613_ at 300 nM were incubated with compounds for 20 min before adding 5 µL CHCHD4-Gst at 100 nM for 2 h. Thirdly, 10 µL of a mixture of 5 µg/mL donor and acceptors beads were incubated for 3 h at 23°C in the dark until reading on EnSpire® multimode plate reader (Perkin Elmer)^17^. For confirmation, best hits (including M30-E05) were validated in triplicate at concentrations of 10 µM and 30 µM.

### Cell lines and patient-derived xenografts (PDX)

Human pediatric osteosarcoma cell lines HOS parental (HOS), HOS-doxorubicin resistant (HOS-R/DOXO), and HOS-methotrexate resistant (HOS-R/MTX), MG63, U20S, 143B, SAOS2 and SAOS2-R/MTX were generously provided by Dr N. Gaspar^19^ and Dr B. Geoerger. Adult cancer cell lines tested which include A549 (non-small cell lung cancer), IGROV-1 (human ovarian carcinoma), IGROV-1 CDDP^R^ (cisplatin-resistant IGROV-1), IGROV-1 VC^R^ (vincristine-resistant IGROV-1), Capan-1 (pancreatic adenocarcinoma), HL-60 (acute promyelocytic leukemia), Jurkat (T-cell leukemia precursor), and NAML6 (acute lymphoblastic leukemia) and the epithelial line of the kidney of a human embryo HEK293 and the proximal tubular cell line derived from normal kidney HK-2 (human kidney 2) were provided by Dr J. Wiels (CNRS UMR 9018, Gustave Roussy, Villejuif). All the cell lines were routinely cultured in Dulbecco’s modified Eagle medium (DMEM, high glucose, GlutaMAX Supplement, Gibco) supplemented with 10% fetal bovine serum (FBS, Sigma-Aldrich, Paraguay Origin) and 1% Penicillin-Streptomycin (Gibco) at 37°C in humidified atmosphere (5% CO_2_ and 95% air), under mycoplasma free conditions. The PDXs GR-OS-9, GR-OS-10, GR-OS-11, GR-OS-12, GR-OS-15, GR-OS-18, GR-OS-20 were established from relapsed or refractory osteosarcoma within the MAPPYACTS PDX project^20^ and kindly provided by Dr B. Geoerger and Dr ME Marques Da Costa. Of note, FBS concentration for *in vitro* PDX secondary culture was 20%.

Human blood samples were obtained from the ‘Etablissement Français du Sang’, Hôpital Saint-Louis, Paris, France and peripheral blood mononuclear cells (PBMCs) were freshly prepared by density gradient centrifugation.

### siRNA transfection

To knock down AIF expression for evaluating the specificity of M30-E05, HOS cells were seeded at densities of 12.500, 18.750, 25.000, 31.250 HOS cells/cm^3^ and transfected with ON-TARGETplus Human *AIFM1* siRNA (Dharmacon, L-011912-00-0005) or ON-TARGETplus GAPDH Control Pool (Dharmacon, D-001830-10-05) for 96h, 72h, 48h, 24h, using 0.2µL, 0.16µL, 0.2µL and 0.08µL of DharmaFECT 1 Transfection Reagent (Dharmacon, T-2001-03), respectively according to the manufacturer’s instructions. For subsequent experiments, 48 hour time point was used. siRNA pool targeting AIF, comprised 4 selected siRNA duplexes each, has a modification pattern that addresses off-target effects caused by both strands (SMART pool).

### Transcriptomic analysis

Under the approval by the steering committee of the MAPPYACTS trial (NCT02613962, Dr B. Geoerger) for 42 refractory or relapsed OS patients and UNICANCER Transbone scientific committee of the OS2006 trial (NCT00470223 – Dr N Gaspar) for 82 diagnostic OS patients, expression of AIF and CHCHD4 was acceded by RNA-sequencing. Mann-Whitney U tests were used to analyze differential expression between diagnostic, non-resistant and resistant OS tumor samples.

### Data availability

Sequencing data and basic clinical annotations from all patients and PDX have been deposited in the BoOSTDataS health data host (https://boostdatasdb.org).

### Gene set enrichment analysis (GSEA)

Cell line RNA-sequencing results were obtained by Dr N. Gaspar and A. Marchais’s team. GSEA was performed using the gseGO function in the clusterProfiler R package, applying the KEGG pathway enrichment to reveal the most significant dysregulated pathways.

## Metabolomics

### a, Sample preparation

**HOS, HOS-R/DOXO, HOS-R/MTX (**300.000 cells per well) were cultured in 6 well-plates and treated with IC_50_ of M30-E05 during 72h. Cells were gently rinsed with cold PBS, and lysed by adding 500 µL of cold methanol/water (9/1, v/v, -20°C) with internal standards (ISTD). After a quick centrifugation, supernatants were pooled, with two wells per condition in a single microtube. Supernatants in microtubes were vortexed at 800 *g* for 5 min and centrifuged at 15000 *g* for 10 min at 4 °C. Supernatants were split in two parts: 150 µL were used for GC-MS experiment in injection vial and 300 µL were used for UHPLC-MS experimentation. LC-MS aliquots were evaporated and dried extracts were resuspended with 150 µL of MilliQ water. Samples were kept at -80°C until injection or transferred in vials for direct analysis by UHPLC/MS.

After evaporation, dry GC-MS aliquot was spiked with 50 µL of methoxyamine (20 mg/mL in pyridine) and stored at room temperature in the dark, overnight. Then, 80 µL of MSTFA was added and final derivatization occurred at 40°C for 30 min. Samples were directly injected into GC-MS.

### b, Targeted analysis of nucleotides and cofactors by ion pairing ultra-high performance liquid chromatography (UHPLC) coupled to a Triple Quadrupole (QQQ) mass spectrometer

Targeted analysis was performed on a RRLC 1290 system (Agilent Technologies) coupled to a Triple Quadrupole 6470 (Agilent Technologies) equipped with an electrospray source.10 μL of sample were injected on a Column Zorbax Eclipse XDB-C18 (100 mm x 2.1 mm particle size 1.8 µm) from Agilent technologies. Gradient mobile phase consisted of water with 2mM of dibutylamine acetate concentrate (DBAA) (phase A) and acetonitrile (phase B). Flow rate was set to 0.4 mL/min, and gradient was performed as follow: initial condition was 90% phase A and 20% phase B, maintained during 3 min. Molecules were then eluted using a gradient from 10% to 95% phase B over 1 min. Column was washed using 95% mobile phase B for 2 min and equilibrated using 10% mobile phase B for 1 min. Scan mode used was the MRM for biological samples. Peak detection and integration of the analytes were performed using the Agilent Mass Hunter quantitative software (B.10.1).

### c, Widely-targeted analysis of intracellular metabolites gas chromatography (GC) coupled to a triple quadrupole (QQQ) mass spectrometer

GC-MS/MS method was performed on a coupling gas chromatography / triple quadrupole 7890B / 7000C (Agilent Technologies, Waldbronn, Germany). The scan mode used was the MRM for biological samples. Peak detection and integration of analytes were performed using the Agilent Mass Hunter quantitative software (B.07.01), exported as tables and processed with R software (version 4.0.3) and the GRMeta package (Github/kroemerlab).

### Western blot

Immunoblotting was performed on cell lysates in NETN buffer supplemented with antiprotease cocktail (ThermoFisher, A32963). Briefly, the cell lysates were separated on 10–15% Tris-glycine SDS–PAGE gel (Invitrogen, NP0322) and transferred onto PVDF membrane. Membranes were blocked in 5% milk in TBS with 0.1% TWEEN-20 (TBST) for 1 hour at room temperature followed by overnight incubation with indicated primary antibodies [MICU1 (1:1000, ATLAS Antibodies, HPA037479), MICU2 (1:1000, ATLAS Antibodies, HPA045511), MCU (1:1000, ATLAS Antibodies, AMAB91189), AIF (1:1000, Cell Signaling, 5318S), CHCHD4 (1:1000, Proteintech, 21090-1-AP), Vinculine (1:1000, Abcam, ab219649), GAPDH (1:1000, Cell Signaling, 2118)] diluted in TBST containing 5% BSA. Membranes were washed and incubated with secondary antibody (Horseradish peroxidase (HRP)-labeled rabbit anti-mouse (1:5000, Jackson ImmunoResearch, 315-035-003) and goat anti-rabbit antibodies (1:5000, Jackson ImmunoResearch, 111-035-144) for 2 hour at room temperature. Bands were visualized using the Substrat HRP Immobilon Western (Millipore, WBKLS0500) and imaged with Amersham™ ImageQuant™ 800. Equal loading was verified by immunoblotting with Vinculin, GAPDH, then normalization of results was performed using free ImageJ software.

### Lactate Deshydrogenase Assay (LDH)

The half maximal inhibitory concentration (IC_50_) at 72 h of M30-E05, Cisplatin, Etoposide for all cell line was determined by CytoTox 96® Non-Radioactive Cytotoxicity Assay, following the manufacturer’s protocol (Promega, G1781). This assay measures the conversion of a tetrazolium (INT) salt to a formazan product, which presents a reddish color. The intensity of color produced is proportional to the number of cells lysed. Absorbance was measured using a 96-well microplate reader at 490nm wavelength (TECAN Infinite M200). The half-maximal inhibitory concentration (IC_50_) was determined using log(agonist) *vs*. response – Variable slope equation (Y=Bottom + (Top-Bottom)/(1+10^((LogEC50-X)*HillSlope))) by GraphPad Prism 10.0 software (GraphPad Software Inc., California, USA).

### Cell death analysis

Cells were seeded in 6-well plates at density of 60.000 cells/well for HOS, and 150.000 cells/wells for all *in vitro* PDX secondary cultures (except 200.000 cells/wells for GR-OS-10 SC and GR-OS-15 SC) and treated with M30-E05 at IC_50_ for 24, 48, 72 h. Culture medium and cells were collected and centrifuged. Supernatant was discarded and cells were stained with APC Annexin V at 12 μg/mL and propidium iodide (PI) at 0.5 mg/mL (BioLegend, 640932). Data acquisition was performed using BD Accuri™ C6 Plus Flow Cytometer. At least 10.000 events were acquired for each sample, and all data were analyzed using FlowJo™ Software (BD Life Sciences).

### Clonogenic assay

HOS cells were seeded at a density of 500 cells/well in 6-well plates. After 24 h, cells were treated with IC_50_ of M30-E05 and kept in the incubator for 10 days under standard conditions. At day 10 after the treatment, colonies were washed twice with PBS and incubated with Crystal Violet (Sigma-Aldrich, HT90132) for 30 min. Images were taken using Amersham™ ImageQuant™ 800 imaging system.

### ATP Assay

ATPlite 1 step Luminescence Assay system from PerkinElmer (Ref.6016736) was used to evaluate the total ATP contents that reflect the catabolic/anabolic status of the cells by measuring total ATP concentration based on the reaction between ATP with added luciferase and D-Luciferine to produce Oxiluciferine and detectable light. Cells were cultured on white 96-well microplates, provided with the kit (PerkinElmer) and total ATP content was measured 24h later with TECAN Infinite 200 Pro microplate reader.

### Bioenergetic profiling using Seahorse XFe96 Assay

To determine the metabolic changes in ATP production, parameters such as the rate of cellular mitochondrial respiration and the rate of extracellular acidification, represented respectively by Oxygen consumption rate (OCR) and Glycolytic efflux rate (GlycoPER), were measured using the Seahorse XFe96 Analyzer (Agilent Technologies) in real-time in 96-well plates. For the untreated conditions, 2.000 cells/well, 4.000 cells/well and 3.000 cells/well were plated 72 h before assay for parental HOS, HOS-R/DOXO and HOS-R/MTX respectively. In the treated conditions, 10.000 cells/ well, 20.000 cells/ well and 15.000 cells/ well were treated during 72 h before assay for parental HOS, HOS-R/DOXO and HOS-R/MTX, respectively. Three different protocols were used to evaluate together or separately OCR and GlycoPER values by injecting different treatments, following manufacturer protocols [Seahorse XF Real-Time ATP Rate Assay (Agilent Technologies, 103592-100); Seahorse XF Cell Mito Stress Test (Agilent Technologies, 103015-100); Seahorse XF Glycolytic Rate Test (Agilent Technologies, 103344-100)].

### Cell counting and normalized analysis

Following the Seahorse assay, cells were incubated with 2.5μg/ml of Hoechst dye in 150μL of PBS solution for 15 min at 37 °C and visualized and counted using Cytation 1 imaging system and Gen5 software (Agilent). Images were captured by DAPI overlay channels or phase-contrast. Raw data analysis and normalization were made using Seahorse analytics (Agilent).

### Confocal microscopy

In a 12 well-plate, glass coverslips were coated with 100µg/mL Poly-D-Lysine at 37°C, overnight. Each coverslip was washed twice with PBS before cell seeding. For the colocalization experiments, HOS cells were seeded at 55.000 cells/3.5cm^2^ well on top of the coated coverslips and allowed to adhere for 24 h. Cells were then exposed to IC_50_ (8.29µM) M30-E05 for 24 or 48h and posteriorly, cells were fixed with 4% PFA for 20 minutes, and permeabilized with 2% Triton X-100 in PBS. After blocking for 40 minutes with 5% BSA in PBS at room temperature, cells were incubated with AIF antibody (1:500, Cell Signaling, 5318) and MTC antibody (1:200, Novus Biologicals, NBP2-32980) in 5% BSA for 45 minutes at room temperature. Cells were then incubated with secondary antibodies Star Orange (1:100, Abberior,), Star Red (1:100, Abberior) and Hoechst (1:10 000, Thermofisher, 62249) in 5% BSA, for 1h at room temperature, protected from light. Abberior mounting media was used to mount the coverslips. For that, the mounting media was warmed in a water bath at 70 °C until it liquidated and was cooled off to room temperature before being used.

To evaluate the mitochondrial network in basal levels, HOS cells were seeded 110.000 cells/3.5cm^2^, HOS-R/DOXO cells were seeded 190.000 cells/3.5cm^2^, HOS-R/MTX cells were seeded 131.000 cells/3.5cm^2^ and allow to adhere for 48h. Fixation and permeabilization were performed as described above. Samples were blocked using 0.5% BSA in PBS for 40 minutes at room temperature. Cells were incubated with Tom20 (1:500, Cell Signaling, 42406) in blocking solution, at 4 °C, overnight, followed by a 1h incubation with the secondary antibody goat anti-rabbit IgG (H+L) Alexa Fluor^TM^ 555 (1:200) (Thermofisher Scientific, A-21428) in blocking solution at room temperature. Coverslips were mounted using FluoroshieldTM with DAPI (Sigma, F6057). 12-bit numerical images were acquired using a confocal Leica SPE DM4000B microscope (pinhole set at 1.0 Airy unit) equipped with a HC PL APO CS2 63x/1.40 oil-immersion objective, using a 405nm, 552nm and 638nm lasers. Images were acquired with zoom adjusted to 1.5 and image size 2048×2048. Z-stacks, with optimal intervals between slides, were acquired for each sample. Mitochondrial network was accessed in Z-stack projection, using Fiji software^21^. To evaluate the colocalization of AIF in mitochondria or in the nucleus, the Z-stacks of each image were analyzed using the JaCoP plugin in Fiji software^21^ ^22^. Pearson coefficient, and Manders’ coefficients M1 and M2, after threshold application, were calculated. Fluorescence intensity profiles of AIF, MTC and Hoechst, following a defined line in x, were evaluated in Z-stack projections of each image. The line was designed to contain the signal of at least three nuclei.

### Transmission electron microscopy

For ultrastructural studies, cells were fixed in 2% glutaraldehyde in 0.1 M Sörensen phosphate buffer (pH 7.3) for 1 h at 4 °C, post-fixed with 2% osmium tetroxide for 1 h at room temperature and stained en bloc in 2% uranyl acetate in 30% methanol for 1 h. Following dehydration through a graded ethanol series, cells were embedded in Epon™ 812. Polymerization was complete after 48 h at 60 °C. Ultrathin sections were stained with standard uranyl acetate and lead citrate and observed with FEI Tecnai 12 electron microscope. Digital images were taken with a SIS MegaviewIII CCD camera. Mitochondria length and width was measured using Fiji software^21^. Ratios length/width closer to 1 indicate that mitochondria are round.

### Mitochondrial membrane potential

To analyze the mitochondrial membrane potential, HOS, HOS-R/DOXO and HOS-R/MTX cells were seeded at 110.000 cells/3.5cm^2^, 190.000 cells/3.5cm^2^, and 131.000cells/3.5cm^2^ , respectively and allowed to adhere for 48h. Cells were then trypsinized and incubated with 100nM Tetramethylrhodamine methyl ester (TMRM) for 10 minutes at 37°C, in the dark. Data acquisition was performed using BD Accuri™ C6 Plus Flow Cytometer. At least 10.000 events were acquired for each sample, and all data were analyzed using FlowJo™ Software (BD Life Sciences).

### Mice studies

All *in vivo* experiments were conducted following the approval of the CEEA26 Ethic Committee (approval number: APAFIS #37810-2022062308201583) agreed by the French Ministry of Research regulations (articles R.214-87 à R.214-126). Animals were purchased at Gustave Roussy (Villejuif, France) and conducted in the respective animal facilities following standard animal regulations, health and care, and ethical controls.

### a, *In vivo* dose escalation

For safety investigation, 25 female NOD.Cg-PrkdcscidIL2rgtm1Wjl/SzJ (NSG) mice were divided into 5 groups (n=5 per group), each receiving different oral doses of the M30-E05 compound: Control (0 mg/kg); 10 mg/kg, 25 mg/kg, 50 mg/kg and 100 mg/kg. M30-E05 was freshly prepared and dissolved in 0.5% methyl cellulose prior to each treatment day. Mice were treated for 4 weeks (5 days per weeks). Behavior, physical appearance, and body weight were monitored and measured to assess any visible potential adverse effects of M30-E05 everyday. At the experiment endpoint, blood and different organs (e.g., liver, lung, kidney, spleen, brain and heart) as well as bone leg were collected at the experimental endpoint, fixed in 4% PFA, and processed for histological analysis. Blood was used for plasma collection.

### b, Subcutaneous patient-derived xenografts (PDX)

2–5mm **s**oft frozen tumor fragments from GR-OS-18 PDX were first xeno-transplanted for amplification and posteriorly transplanted subcutaneously into 35 immunocompromised NOD.Cg-PrkdcscidIL2rgtm1Wjl/SzJ (NSG) 35 mice (21 females and 14 males) under anesthesia (3% induction, 2% maintenance isoflurane, and 1.5 L/min air). 17 mice were assigned to the control group, receiving the vehicle only for 5 days per week for two weeks, while 18 mice were assigned to the treatment group, receiving 100 mg/kg M30-E05 on the same schedule. Clinical observations included monitoring skin changes, behavior, posture, responses to handling, and abnormal movements. Body weight and tumor volume were measured twice weekly. Tumor volume was calculated according to the equation: V (mm^3^) =width2 (mm^2^) ×length (mm)/2. The experiments lasted until tumors reached specific endpoints detailed in the ethical projects. Upon euthanasia, blood, tumors and various organs (bone, liver, lung, kidney, spleen, and brain) were collected for mass spectrometry and western-blot (snap-frozen in liquid nitrogen) and histology analysis. Tissues fixed in 4% PFA and included in FFPE blocs were sectioned and stained with hematoxylin and eosin and analyzed by an experienced histopathologist.

### ELISA

Blood from 25 treated mice and control (vehicle-treated) mice used for the *in vivo* efficiency were centrifuged at 2000 rpm, 5 min at 4°C for plasma collection. The mouse cardiac troponin-I (TNNI3) levels in plasma were measured using an ELISA Kit (Invitrogen, EEL112), following the manufacturer’s instructions.

### Histology and immunohistochemistry

Organs were fixed in 4% PFA and embedded in paraffin; 4μm sections were stained with hematoxylin-eosin-safranin for morphology. Paraffin sections were processed for heat-induced antigen retrieval (ER2 corresponding EDTA buffer pH9) for 20 min at 100°C. Slides were incubated with a mouse monoclonal anti-human Ki67 antibody (1:20, Agilent Dako, clone MIB1) for 1h at room temperature. The nuclear signal was revealed with the Klear mouse kit (GBI labs). Single representative whole tissue section from each animal was digitized using a slide scanner NanoZoomer 2.0-HT (Hamamatsu Photonics, C9600-13).

### Multiplex immunofluorescence analysis

The immunofluorescence multiplex panel was performed using sequential indirect immunolabeling with tyramid signal amplification technique. After tissue dewaxing at 72 °C during 30 sec, an antigen retrieval step at pH 9, at 100 °C for 20 min was done with an epitope retrieval solution (ER2). Next, endogenous peroxydases and non-specific binding sites were blocked with hydrogen peroxide (10 min at room temperature) and PKI blocking solution (5 min at room temperature, without wash) prior to the application of the primary antibody for 30 min at RT. This one was then recognized by an anti-rabbit HRP-polymer conjugated secondary antibodies or by an ultrapolymer HRP donkey anti-goat incubated for 30 min at RT. The formation of highly reactive fluorochrome-tyramid complex (Akoya Opal®), able to bind the tyrosin of tissue in the vicinity of the antigen-antibody complex, was then catalyzed by HRP during an incubation step of 20 min at RT. The antibodies complex (but not the tyramids) were thereafter removed during a stripping step by incubation in ER solution at pH 6, at 97 °C during 20 min, before the next immuno-labeling cycle. Slides were counterstained spectral DAPI (Akoya Biosciences) was added to the slides for 5 min and coverslips were mounted with mounting medium ProLongTM Diamond Antifade Mountant (Invitrogen^TM^, ThermoFisher Scientific).

Multiplex labeling was performed on an automated staining system Leica BOND RX (Leica Biosystems, Nussloch, Germany). The primaries antibodies: anti-MTC (Novus Biologicals, NBP2-32980, clone 113-1), anti-AIF (Abcam, ab288370, clone 4B2); anti-CHCHD4 (Proteintech, 21090-1-AP, polyclonal); anti-MICU1 (Abcam, ab190114, goat polyclonal). Tissue sections were sequentially incubated with the following reagents and conditions (dilutions, OPAL) in this order: anti-MTC 1:200, Opal® 620 1:100; anti-AIF 1:500, Opal® 520 1:400; anti-CHCHD4 1:1000, Opal® 520 1:200. Slides were scanned with a Vectra Polaris for analysis.

### Computational study

The human AIF monomer three-dimensional structure (PDB ID: 1M6I co-crystalized with one flavin adenine dinucleotide (FAD) was used^23^. Missing residues of the monomer structure (except those at terminal ends) were first added using SWISS-MODEL webserver^24^. AIF structure was submitted to a 500 ns molecular dynamics (MD) simulation in explicit solvent, using GROMACS software^25^ with AMBER99SB-ILDN^26^ and TIP3P^27^ all-atom force field. Then, the 5 most populated clusters of the protein conformational ensemble were extracted from the MD trajectory and used as receptors in subsequent docking calculations.

Each studied ligand was blindly docked 100 times on the whole surface of each of the AIF clusters (without specifying any preferred binding site), using AutoDock Vina^28^. Each docking calculation generating 10 binding modes, a total of 5.000 binding modes of each ligand was obtained on AIF protein. In addition, the docking results of M30-E05 on human AIF were compared to those obtained on murine AIF (PDB ID: 1GV4)^29^ using a similar protocol.

### Statistics

Statistical analysis was performed using GraphPad Prism 10.0. Mann-Whitney U test was applied for RNA-Seq analysis, tumor volume curves and IHC analysis of AIF, Ki-67 expression in tumor: p<0.033 (*), p<0.002 (**), p<0.001 (***). One-way ANOVA test was used for statistical analysis of biological experiments: p<0.05 (*), p<0.01 (**), p<0.001 (***), p<0.0001 (****).

## Results

### 1. Identification and synthesis of M30-E05 as an AIF/CHCHD4 inhibitor

AIF and CHCHD4 associate in a protein-protein complex that is necessary for the biogenesis of respiratory chain complexes and cell survival^18^. A robotized high throughput screening (HTS) assay to identify compounds interrupting the interaction between AIF and CHCHD4 has been developed using the Alphascreen technology^17^ (Fig. 1a). To that end, we produced tagged HIS-AIF_103-613_ and GST-CHCHD4 recombinant proteins in *E. coli*^17^ and purified them to homogeneity as shown by Western blot (Fig. 1b). Then, taking advantage of their tags, we coupled them with donor and receptor alpha beads and determined the optimal concentrations of proteins to measure their interaction by light emission in 384-well plates (Revvity) (Fig. 1c). As a positive control of protein-protein interaction inhibitor (PPi) capable to fully interrupt the AIF/CHCHD4 interaction, a synthetic peptide, called N27, corresponding to the N-terminal 27 amino acids of CHCHD4 was used^18^.

**Figure 1.**
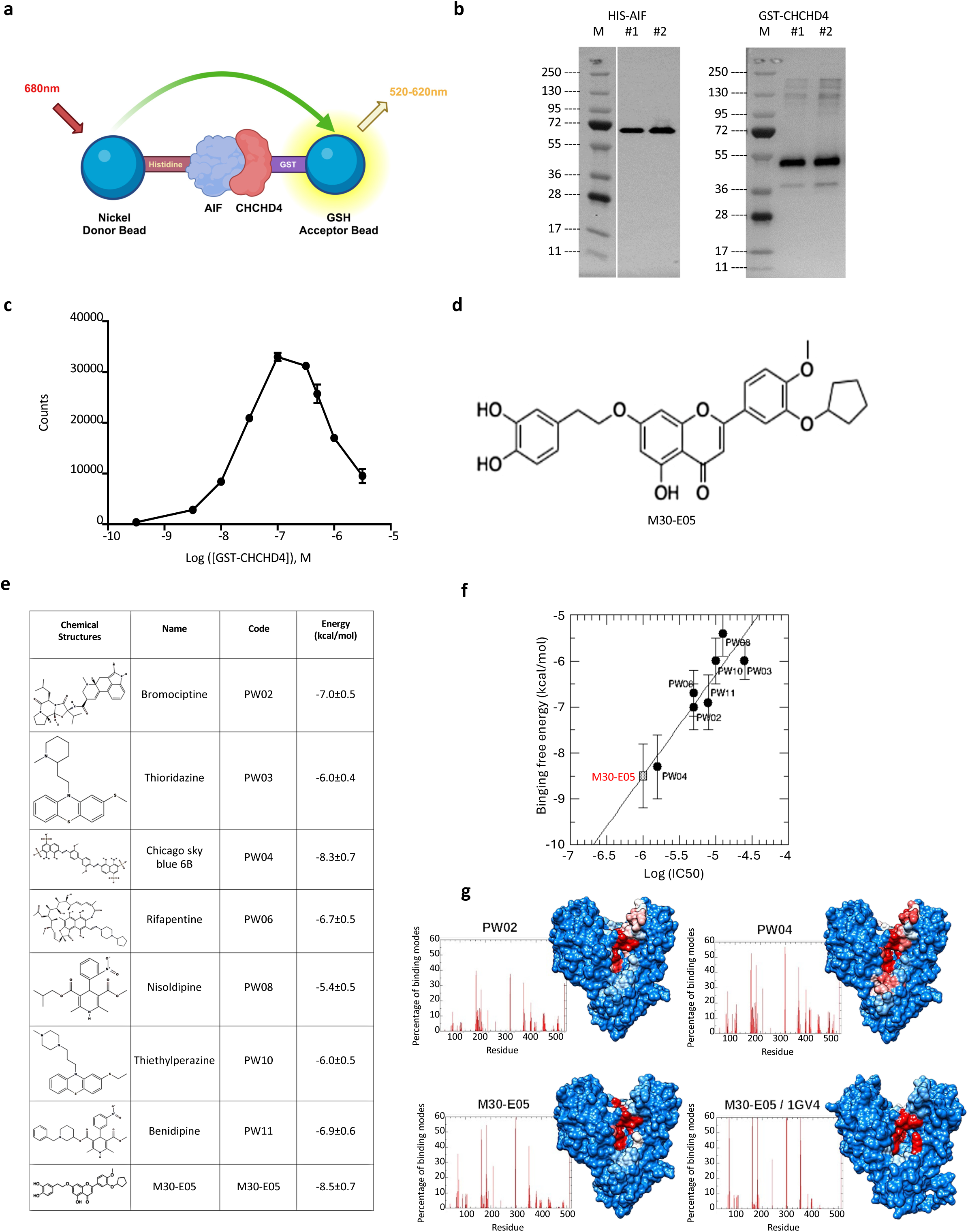
High throughput screening assay for the robotized screening of PPi of AIF-CHCHD4 complex and M30-E05 synthesis. **a**, Scheme of the *in vitro* Alpha screening assay to measure the interaction between HIS-AIF and GST-CHCHD4 coupled to donor and acceptor beads. Adapted from manufacturer’s protocol **b,** Human AIF_103-613_ and CHCHD4 were produced as recombinant proteins in *E. coli* and their purity was analyzed by SDS-PAGE and western-blot. #, independent production and purification experiments. **c,** Optimization of the ratio AIF/CHCHD4 coated on beads for the detection of their interaction. **d,** Chemical formula of M30-E05. PPi, protein-protein interaction inhibitor. **e,** Chemical structure of experimentally identified inhibitors of AIF/CHCHD4 interaction and binding free energy for human AIF estimated by Autodock Vina docking calculations. PW, Prestwick. **f,** Correlation between AutoDock Vina binding free energies of 7 Prestwick compounds (black circle) on human AIF and their log(IC_50_) experimentally measured. The red square on the correlation line indicates M30-E05 binding free energy. **g,** Contact frequency of AIF residues with PW02, PW04, and M30-E05 binding modes (graphs) and localization of AIF surface residues most frequently in contact (blue, white and red colors indicate residues that are contacted less than 10%, between 10% and 20%, and more than 20% out of all the ligand binding modes, respectively). The bottom-right graph and structure are the results of M30-E05 docking on murine AIF.

In a pilot experiment, the miniaturization and robotization of assay was validated by a z’ factor comprised between 0.5 and 1. Subsequently, we conducted a screening campaign using the Prestwick library of 1.280 FDA-approved molecules, a library of 2.901 new compounds from the French National Chemical library, and 99 newly synthesized molecules from the BioCIS laboratory. Compounds (tested at 10 µM) were evaluated for their capacity to reduce the alpha signal, compared to N27 as a positive control. Consequently, we identified 11 hits coming from the Prestwick library^17^ and 4 from the BioCIS laboratories. Among these, M30-E05 interrupted the AIF/CHCHD4 complex and presented the best activity *in vitro* and *in cellulo*, we decided to pursue the characterization of its bioactivity in an anti-cancer therapeutic perspective.

Next, M30-E05 was resynthesized through a 7-step strategy at multigram scale (Fig. 1d, Supplementary Note, patent request EP24202518.7). The synthesis started from diosmetin (**1**), which was selectively alkylated in 3 steps with a cyclopentyl group at C-3’ yielding flavone (**2)**. A benzyl protection of the most reactive C-7 hydroxygroup was necessary to achieve this synthesis. In parallel, the synthesis of iodide (**4**) was achieved by protection of catechol (**3**) as an orthoester, followed by a Finkelstein reaction. The alkylation of the most reactive C-7 hydroxygroup of (**2**) with (**4**) yielded the expected coupling product. Subsequent hydrogenation with catalytic HCl, directly yielded M30-E05, by reducing selectively the aromatic ketone and deprotecting the orthoester function simultaneously.

### 2. *In silico* characterization of M30-E05 binding free energy and binding site on AIF

We first used the docking procedure described in the Materials and Methods section on 7 compounds from the Prestwick chemical library (Fig. 1e) for which an experimental IC_50_ of the AIF/CHCHD4 interactions could be measured by the Alphascreen technology^17^. As shown in Fig. 1f, ligand average binding energies yielded by AutoDock Vina on AIF monomer could be fairly correlated with the experimental log(IC_50_), indicating that our computational method is a helpful tool for identifying potential efficient inhibitors of AIF/CHCHD4 interactions.

In a second step, the M30-E05 compound identified from the BioCis Library (Fig. 1e), which could also inhibit the AIF/CHCHD4 complex, was docked on the AIF monomer using the same protocol. Our calculations show that M30-E05 has a better binding free energy of -8.5±0.7 kcal/mol, which is better than another 7 Prestwick compounds (Fig. 1f).

To gain insight into the mechanism of action of these AIF/CHCHD4 inhibitors, we further investigate the preferred binding site of the two best hits from Prestwick, PW02, PW04, and M30-E05 molecules on AIF monomer. Strikingly, as shown in Fig. 1g, these molecules preferentially bind the same surface area on AIF protein, which corresponds to the NAD binding pocket. This strongly suggests that these molecules might competitively inhibit NAD binding, which in turn would prevent AIF dimerization.

Finally, we performed a similar docking procedure of M30-E05 on the murine AIF structure (PDB ID: 1GV4). The obtained average binding free energy is -9.0 ± 0.7 kcal/mol which is slightly lower than that of the human protein. Furthermore, the preferential binding site of M30-E05 on murine AIF is also found to be the NAD binding pocket (Fig. 1g). Overall, M30-E05 binds similarly the murine and human AIF protein.

### 3. M30-E05 reduces viability of cancer cell lines and impacts mitochondria structure and function

Osteosarcoma was selected as a relevant cancer model expressing significant levels of AIF and CHCHD4 to test the activity of M30-E05 based on RNA-seq transcriptomic analysis of patient cohorts from MAPPYACTS^30^ and OS 2006^31^ studies, which include patients at relapse and diagnosis respectively (Supplementary Fig. 1 a,b). None of the osteosarcoma patients harbored mutations of the *AIFM1* gene, although 2 patients /628 patients of MAPPYACTS cohort (one Ewing sarcoma patient and one brain central nervous system patient) presented somatic *AIFM1* gene mutations^30^. Accordingly, several osteosarcoma PDX demonstrated significant expression of AIF and CHCHD4 (Supplementary Fig. 1c).

To assess the activity of M30-E05, a panel of osteosarcoma cell lines, including both parental and drug-resistant cell lines (HOS, HOS-R/DOXO, HOS-R/MTX, MG63, U2OS, 143B, SAOS2 and SAOS-R/MTX) was treated with M30-E05 (8 doses, ranging from 1 to 100 μM dissolved in 0.002% DMSO) for 72h. Cell viability was then evaluated using a lactate dehydrogenase (LDH) assay, and the IC_50_ determined using GraphPad Prism software. IC_50_ for HOS cells was 8.29 μM with a total cell viability loss and across all cell lines, the IC_50_ was approximately 10 μM, independent of resistance to doxorubicin and methotrexate (Fig. 2a). Since doxorubicin resistance is mediated by increased PgP efflux activity, the obtained IC_50_ suggests that M30-E05 is not a PgP substrate.

**Figure 2.**
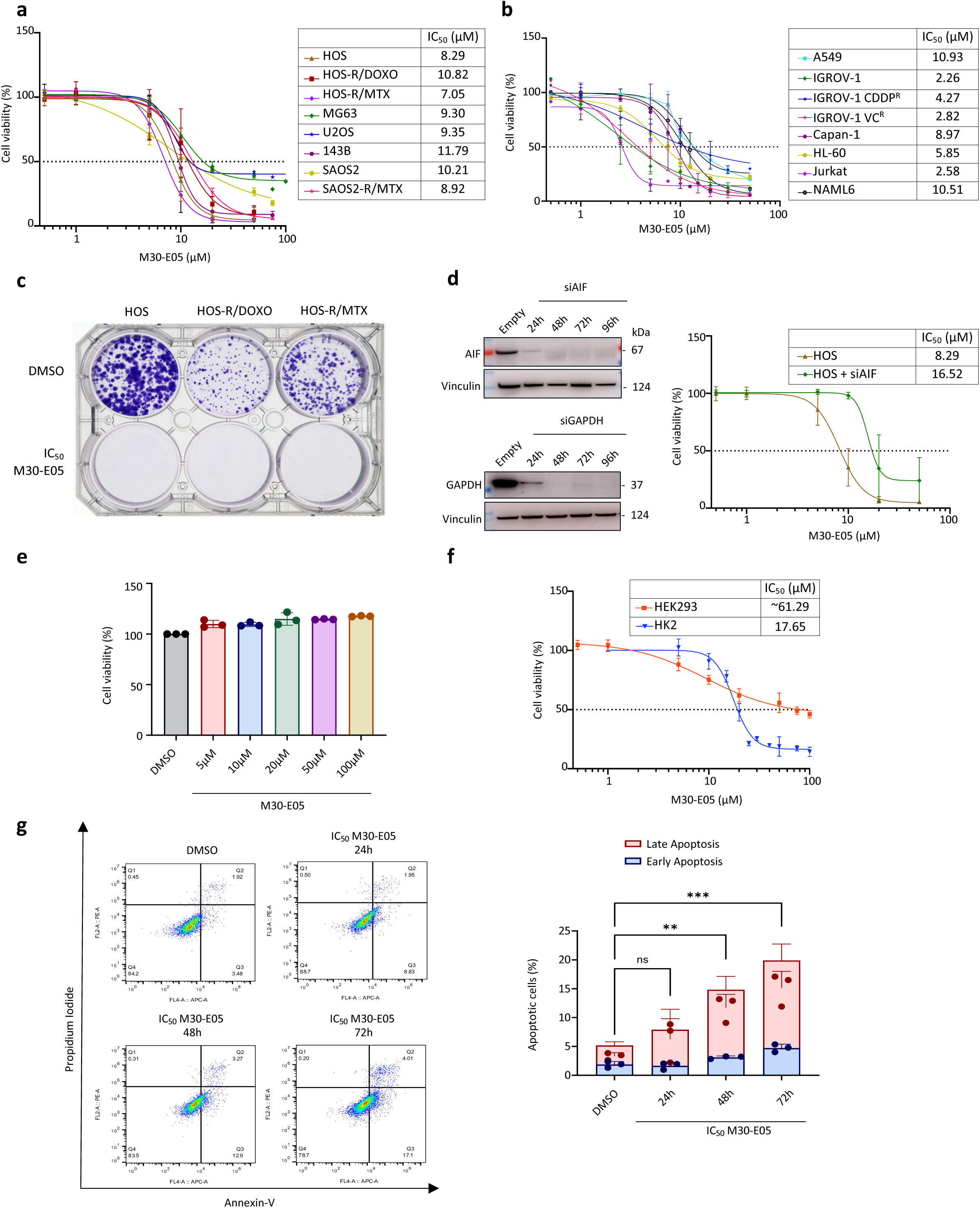
Cytotoxic effects analysis of M30-E05 on cancer cell lines and non-cancerous models. **a,** Cell viability of 8 pediatric osteosarcoma cell lines treated with M30-E05. Dose-response curves represent cell viability measured using a LDH assay at increasing concentrations of M30-E05 after 72 hours of treatment. Individual experiments are shown (n > 2). IC_50_, inhibitory concentration yielding to 50% cell viability. 0.002% DMSO was used as the solvent of M30-E05 and as a control for calculation of percentage of viability. **b,** Cell viability of 10 adult cancer cell lines treated with M30-E05. Dose-response curves represent cell viability measured using a LDH assay at increasing concentrations of M30-E05 after 72 hours of treatment. Individual experiments are shown (n > 2). 0.002% DMSO was used as the solvent of M30-E05 and as a control for calculation of percentage of viability. **c,** Image of a 6-well plate showing fixed colonies subjected to crystal violet staining of HOS, HOS-R/DOXO, and HOS-R/MTX cells after 10 days of culture, treated with either 0.002% DMSO as a negative control or IC_50_ M30-E05. **d,** Representative Western blot analysis of AIF-siRNA (siAIF) treatment over time (top panel), with siGAPDH as a positive control (bottom panel). Based on this, AIF was silenced 48h before exposure to increasing concentrations of M30-E05. Dose-response curves were generated after 72h of treatment. Vinculin was used as a loading control. Experiment has been repeated three times. **e,** Cell viability of peripheral blood mononuclear cells (PBMCs) isolated from human donors treated with either 0.002% DMSO as a negative control or with M30-E05 at various concentrations (5 µM, 10 µM, 20 µM, 50 µM, and 100 µM), assessed via LDH assay after 72 hours. Individual experiments are shown (n = 3). **f,** Cell viability of non-cancerous cell lines treated with M30-E05, measured using an LDH assay. Dose-response curves show cell viability at increasing M30-E05 concentrations after 72 hours of treatment. Individual experiments are shown (n > 2). Non-cancerous cell lines tested include HEK293 (immortalized human embryonic kidney cells) and HK2 (immortalized normal adult human kidney cells). **g,** Annexin V/PI double-staining assay of HOS cells showing flow cytometric analysis of IC_50_ (8.3 µM) M30-E05-induced apoptosis at different time points (24h, 48h, and 72h). 0.002% DMSO was used to treat HOS cells and considered as a negative control. Statistical graph of Annexin V-FITC/PI staining results, with data representing the mean of three independent experiments. Annexin V+/PI− cells are early apoptotic, and Annexin V+/PI+ cells are late apoptotic. Statistical significance was assessed using one-way ANOVA with Šídák’s multiple comparisons test. Significance levels are indicated as follows: not significant (ns), p < 0.0332 (*), p < 0.0021 (**), p < 0.0002 (***), and p < 0.0001 (****).

M30-E05 also reduced the viability of several tumor cell lines derived from ovarian cancer (both parental and drug-resistant cell lines), lung cancer, pancreatic cancer, as well as leukemia (Fig. 2b). Notably, M30-E05 exhibited an IC_50_ of 2-3 μM in ovarian cancer cell lines and some leukemia cell lines.

Further characterizations were performed using 3 human osteosarcoma cell lines, HOS, the parental cell line, and its 2 drug-resistant derivatives, HOS-R/DOXO and HOS-R/MTX, obtained by culture in sublethal doses of doxorubicin (DOXO) and methotrexate (MTX)^19^. The three cell lines expressed immunoblot-detectable AIF and CHCHD4 (Supplementary Fig. 1d). Immunofluorescence and electron microscopy revealed that, as compared to HOS, HOS-R/DOXO and HOS-R/MTX cells showed a fragmented mitochondrial network (Supplementary Fig. 2 a,b). Moreover, as compared to HOS, HOS-R/DOXO cells exhibited a reduction in mitochondrial transmembrane potential, mitochondrial oxygen consumption rate (OCR), and total ATP content (Supplementary Fig. 2 c,d,e). HOS-R/MTX displayed close-to-normal mitochondrial potential and respiration rate, but a subnormal ATP content (Supplementary Fig. 2c,d,e). Both resistant cell lines manifested a shift in metabolites abundance towards lipid and amino acid metabolism compared to the parental HOS cell line (Supplementary Fig. 2f,g).

In a clonogenic assay conducted on the three osteosarcoma cell lines, all cells undistinguishably succumbed to treatment with IC_50_ M30-E05 of each cell line within 10 days (Fig. 2c). However, M30-E05 was less effective against HOS cells subjected to AIF knockdown by transfection with a suitable siRNA (IC_50_ = 16.52 μM) (Fig. 2d). Importantly, M30-E05 at 100 µM failed to affect the viability of freshly isolated human peripheral blood mononuclear cells (PBMCs) (Fig. 2e), while its IC_50_ of M30-E05 for non-cancerous HEK293 and HK2 cells were ∼62 μM and 18 μM, respectively (Fig. 2f). When treated at its IC_50_ (8.29 µM), M30-E05 induced HOS cell apoptosis as determined using Annexin V/PI staining and cytofluorometric analyses over several days (Fig. 2g).

Of note, the immunoblot-detectable protein expression of AIF, CHCHD4 and two substrates of the AIF/CHCHD4 pathway, MICU1 and COX17, revealed an early depletion of all these proteins within 6h of treatment with the IC_50_ (8.29 µM) of M30-E05 in HOS cells (Fig. 3a). Expression of VDAC, a mitochondrial outer membrane protein independent of AIF/CHCHD4, remains unaffected under these conditions. Confocal immunofluorescence microscopy revealed that M30-E05 induced the fragmentation of the mitochondrial network and the relocalization of mitochondria towards the nuclear envelope without any chromatin condensation at 24h and 48h (Fig. 3b). AIF remained localized at mitochondria and failed to translocate to nuclei (Fig. 3c). Transmission electron microscopy confirmed morphological alterations of mitochondria, including increased matrix density and cristae loss in M30-E05-treated cells (Fig. 3d).

**Figure 3.**
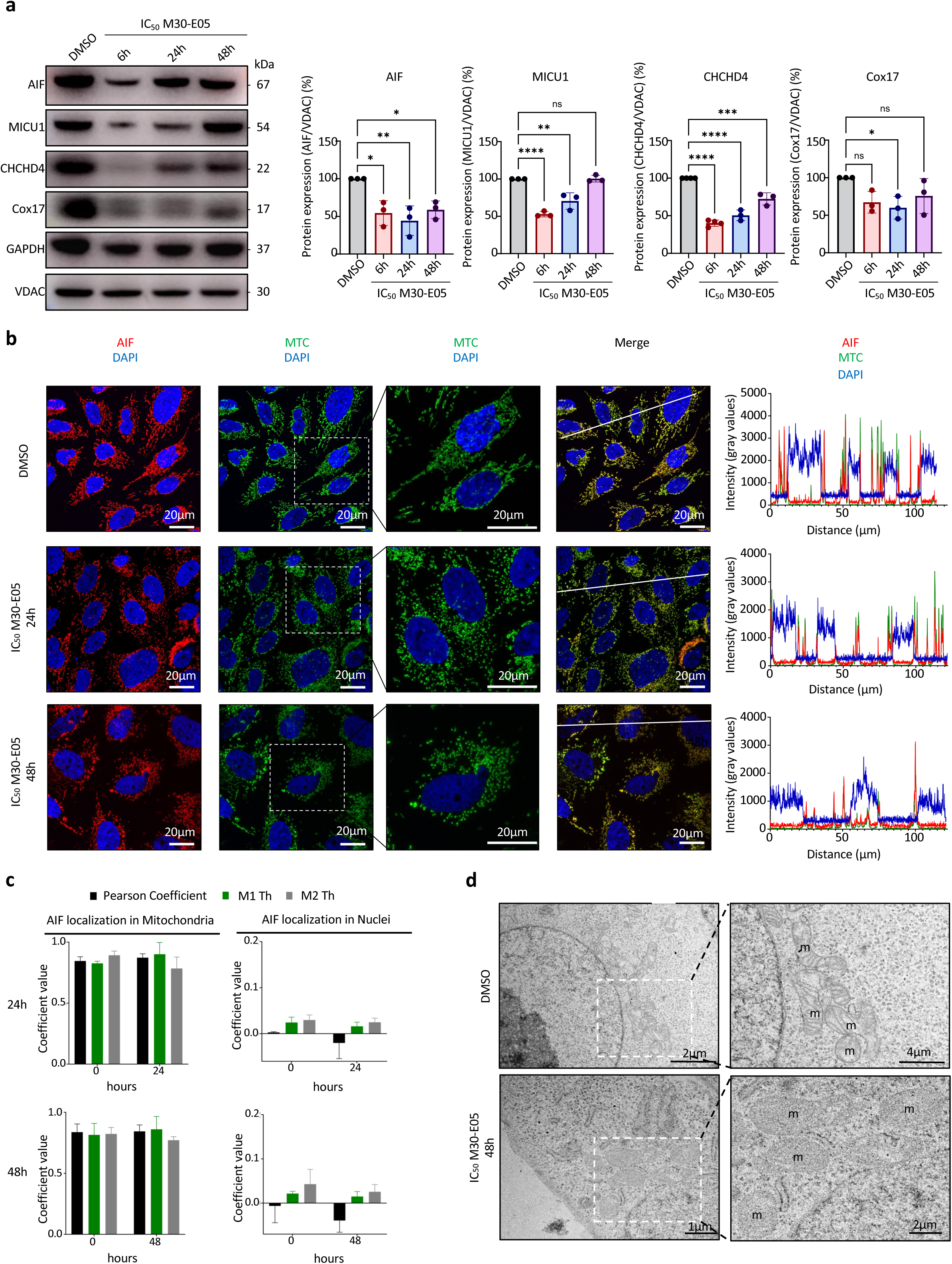
Effects of M30-E05 on mitochondrial AIF/CHCHD4 complex, network integrity, and structural morphology in HOS cell line. **a,** Western Blot analysis showing the expression levels of AIF, CHCHD4 and their substrates Cox17 and MICU1 in response to either 0.002% DMSO as a negative control or IC_50_ (8.3 µM) of M30-E05 treatment over time (6h, 24 h, and 48h). Vinculin and VDAC were used as loading controls. Data are presented as the mean ± SD from four independent experiments (n = 4). Statistical significance was assessed using ANOVA with Sidak’s correction for multiple comparisons. Significance levels are indicated as follows: not significant (ns), p < 0.0332 (*), p < 0.0021 (**), p < 0.0002 (***), and p < 0.0001 (****). **b,** Representative confocal images of HOS cells exposed to either 0.002% DMSO as a negative control or IC_50_ (8.3 µM) M30-E05 for 0, 24 and 48h. AIF colocalization in mitochondria labelled with MTC and in the nucleus stained with DAPI is evaluated. Intensity profiles of AIF, MTC and DAPI, following a defined line in X, and its superposition is analyzed in Z-stack projections of each condition and are represented in graphs on the right. The line was chosen to contain the signal of at least three nuclei. Four independent experiments were performed. **c,** The Pearson Coefficient and Manders’ coefficients M1 and M2, after threshold application were calculated in each image, from four independent experiments, using the JaCoP plugin in Fiji Software. **d,** Representative transmission electron microscopy images of HOS cells treated with either 0.002% DMSO as a negative control or with IC_50_ (8.3 µM) M30-E05 for 48 h, showing that the compound induced cristolysis and changes in mitochondrial ultrastructure. m: mitochondrion

Concomitantly, M30-E05 significantly decreased OCR in HOS cells starting at 6h post-treatment, with a maximal effect at 24h (Fig. 4a). Despite impaired mitochondrial respiration, total cellular ATP levels increased (Fig. 4b), suggesting an increase in ATP production by glycolysis and/or a reduction of ATP consumption.

**Figure 4.**
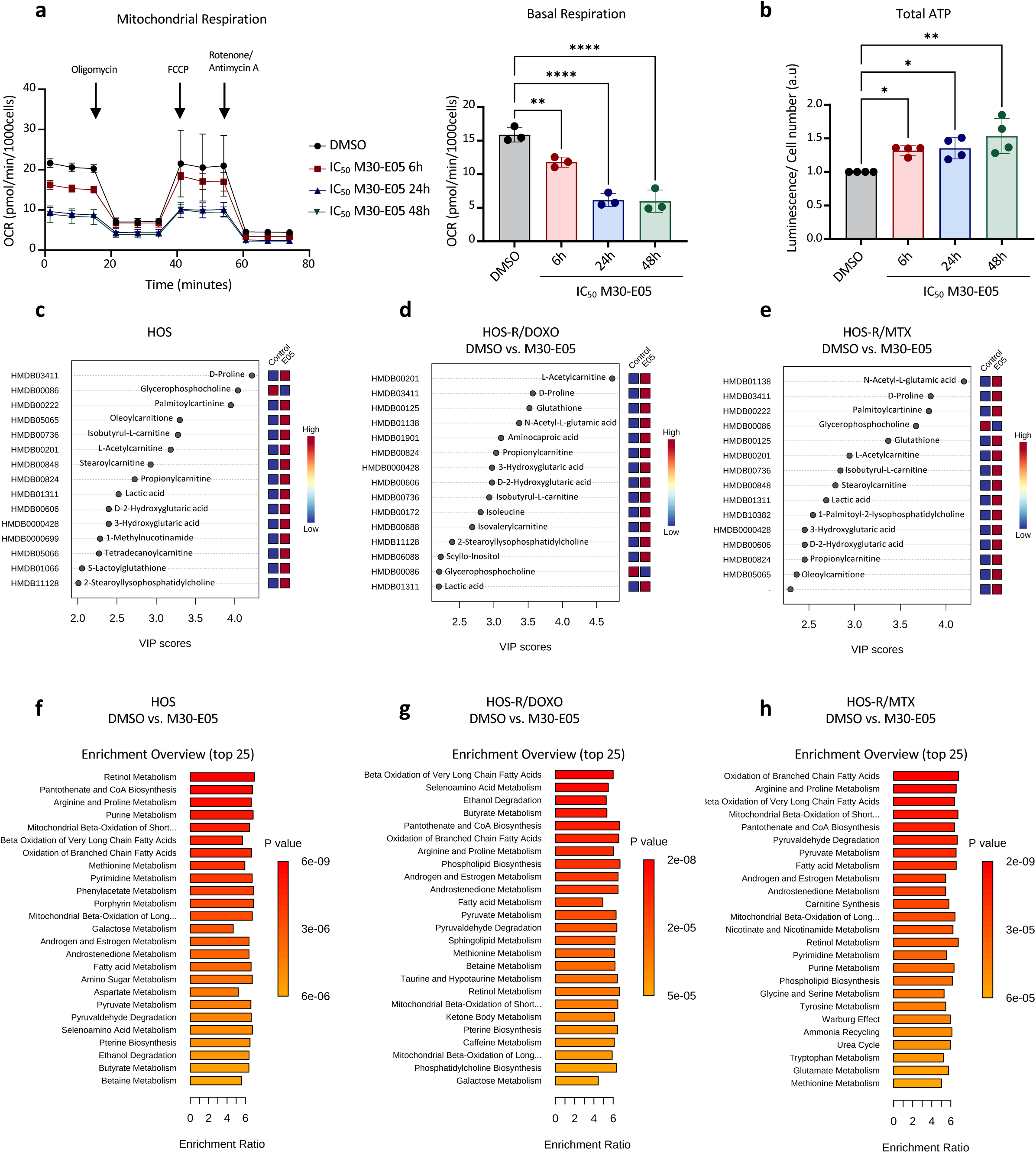
Effects of M30-E05 on energetic metabolism and metabolic reprogramming in osteosarcoma cell lines. **a,** Seahorse XFe96 Mito Stress Test graph displaying the oxygen consumption rate (OCR; pmol/min/1.000 cells) over time in HOS cells pre-treated with either 0.002% DMSO as a negative control or with IC_50_ (8.3 µM) M30-E05 for 6, 24, and 48h (left). The test was conducted using the following compounds: Oligomycin (2.5 µM), FCCP (1 µM), and Rotenone/Antimycin A (0.5 µM). Basal respiration values were extrapolated from the kinetic graph. OCR measurements were normalized to cell counts determined by nuclei DAPI staining. Data represent the mean ± SD of three independent experiments, each with at least six technical replicates. Statistical analysis was performed using ANOVA with Sidak’s correction for multiple comparisons, with significance levels indicated as follows: ns, not significant; p < 0.0332 (*); p < 0.0021 (**); p < 0.0002 (***);p < 0.0001 (****). **b,** ATP levels were semi-quantified over time in HOS cells pre-treated with either 0.002% DMSO as a negative control or with IC_50_ (8.3 µM) M30-E05 for 6, 24, and 48 h, using the ATPlite assay. **c, d, e,** VIP (Variable Importance in Projection) score plot depicting metabolites that significantly differentiate between 0.002% DMSO as a negative control and IC_50_ (8.3 µM) M30-E05-treated osteosarcoma cell lines following 72h of treatment. Two vertical box plots on the right illustrate metabolic variations in 0.002% DMSO *versus* M30-E05-treated c) HOS, d) HOS-R/DOXO, e) HOS-R/MTX, respectively. **f, g, h,** Enrichment analysis of metabolites that differ significantly between 0.002% DMSO as a negative control and IC_50_ (8.3 mM) M30-E05-treated cells in (f) HOS, (g) HOS-R/DOXO, and (h) HOS-R/MTX cell lines following 72h of treatment. The enrichment ratio represents the observed number of metabolites in a specific metabolic pathway divided by the expected number. Metabolic pathways are ranked by p-value, with the most significant pathways at the top.

Mass spectrometric metabolomics was performed to further elucidate the M30-E05 effects on the metabolic profiles of 3 HOS cell lines (parental and resistant) (Fig. 4 c-h). The results highlighted significant impact of M30-E05 on mitochondrial β-oxidation of short-and long-chain fatty acids, as well as metabolic pathways affecting amino acids including arginine and proline (Fig. 4 c-h).

### 4. M30-E05 effects on secondary cultures of osteosarcoma PDX

Patient-derived xenografts (PDX) reflect the complexity of heterogeneity of human cancers more accurately than cell lines. After a first passage through immunodeficient mice, PDX can be either transplanted again into mice or subjected to secondary culture *in vitro*. Usually, even osteosarcoma PDX that grow *ex vivo* cannot be cultured >3 passages without losing their fitness^20^. PDX from 3 patients that were xenografted in NSG mice either subcutaneously (SC) or orthotopically/paratibially (PT) all expressed AIF and CHCHD4 at the mRNA and protein levels (Fig. 5a,b) with a clear mitochondrial localization of AIF and CHCHD4 (Supplementary Fig. 3). Transmission electronic microscopy observation revealed some variability in the morphology (round *vs* elongated) and ultrastructure of the mitochondria that is in line with the variability in basal OCR of the PDX and their grafting location, suggesting an effect of the tumor microenvironment (Fig. 5d, Supplementary Fig. 4). Secondary subcultures of these PDX models showed reduced viability upon M30-E05 treatment, with IC_50_ values ranging from 17 to 40 µM (Fig. 5c), which is comparable to IC_50_ obtained with etoposide and cisplatin, two standard chemotherapeutic agents used in clinics (Supplementary Fig. 3a,b). M30-E05 treatment also tended to decrease basal OCR in all PDX models (Fig. 5d) and accompanied by a net decrease in AIF, MICU1 and CHCHD4 protein levels in some PDX cultures (Fig. 5e).

**Figure 5.**
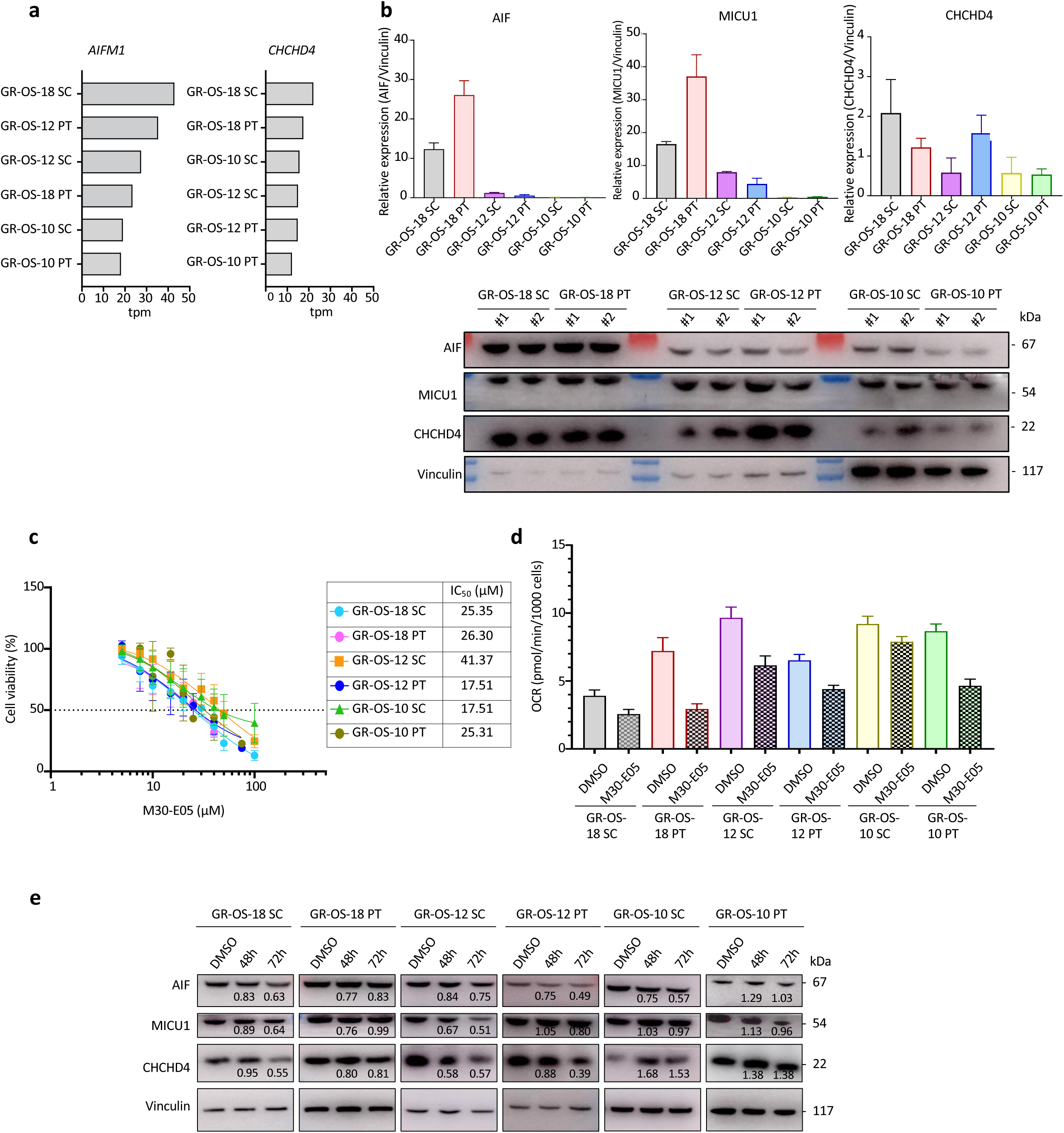
Characterization of AIF/CHCHD4 complex and cellular responses in secondary cultured in vitro PDX following M30-E05 treatment. **a,** Normalized gene expression levels of *AIFM1* and *CHCHD4* from RNA-seq data across six different PDX samples, ranked from highest to lowest expression. Tpm, transcripts per million. **b,** Representative Western blot analysis showing the expression levels of the AIF, CHCHD4 and its substrates (MICU1) across six different secondary cultured *in vitro* PDX samples with 2 biological replicates (#1 and #2 for each PDX). Vinculin was used as a loading control. Experiments has been repeated twice. **c,** Cell viability of 6 different secondary cultured *in vitro* PDX samples treated with different concentrations of M30-E05, was measured using an LDH assay. Dose-response curves represent cell viability at increasing concentrations of M30-E05 after 72h of treatment. Individual replicates are shown (n > 2). **d,** Basal respiration displaying the oxygen consumption rate (OCR; pmol/min/1.000 cells) in 6 different secondary cultured *in vitro* PDX samples pre-treated with either 0.04% DMSO as a negative control or with 20µM M30-E05 for 24 h. Seahorse Xfe96 MitoStress Test was conducted using the following compounds: Oligomycin (2.5 µM), FCCP (1 µM), and Rotenone/Antimycin A (0.5 µM). Basal respiration values were extrapolated from the kinetic graph. OCR measurements were normalized to cell counts determined by DAPI nuclei staining. Data represent the mean ± SD of one to two independent experiments, each with at least six technical replicates. **e,** Representative western blot showing the expression levels of the AIF, CHCHD4 and its substrate (MICU1) in response to either 0.04% DMSO as a negative control or 20 µM M30-E05 treatment over time (48h and 72h). Vinculin was used as a loading control.

### 5. Safety and antitumor activity of M30-E05 in subcutaneous osteosarcoma PDX *in vivo*

As a preclinical safety assessment, a dose-escalation study of M30-E05 was performed in NSG mice. M30-E05 was administered orally for 5 days per week over 4 weeks at doses ranging from 10 mg/kg up to 100 mg/kg, using 0.5% methylcellulose as a control (vehicle) (Fig. 6a). Throughout this period, the behavior, fur aspect, body weight (Fig. 6a) and circulating troponin (TNNI3) levels were not affected (Fig. 6b). Moreover, Hematoxylin/Eosin staining analysis of major organs (heart, lung, liver, kidney, spleen, brain) revealed no morphological abnormalities (Fig. 6c).

**Figure 6.**
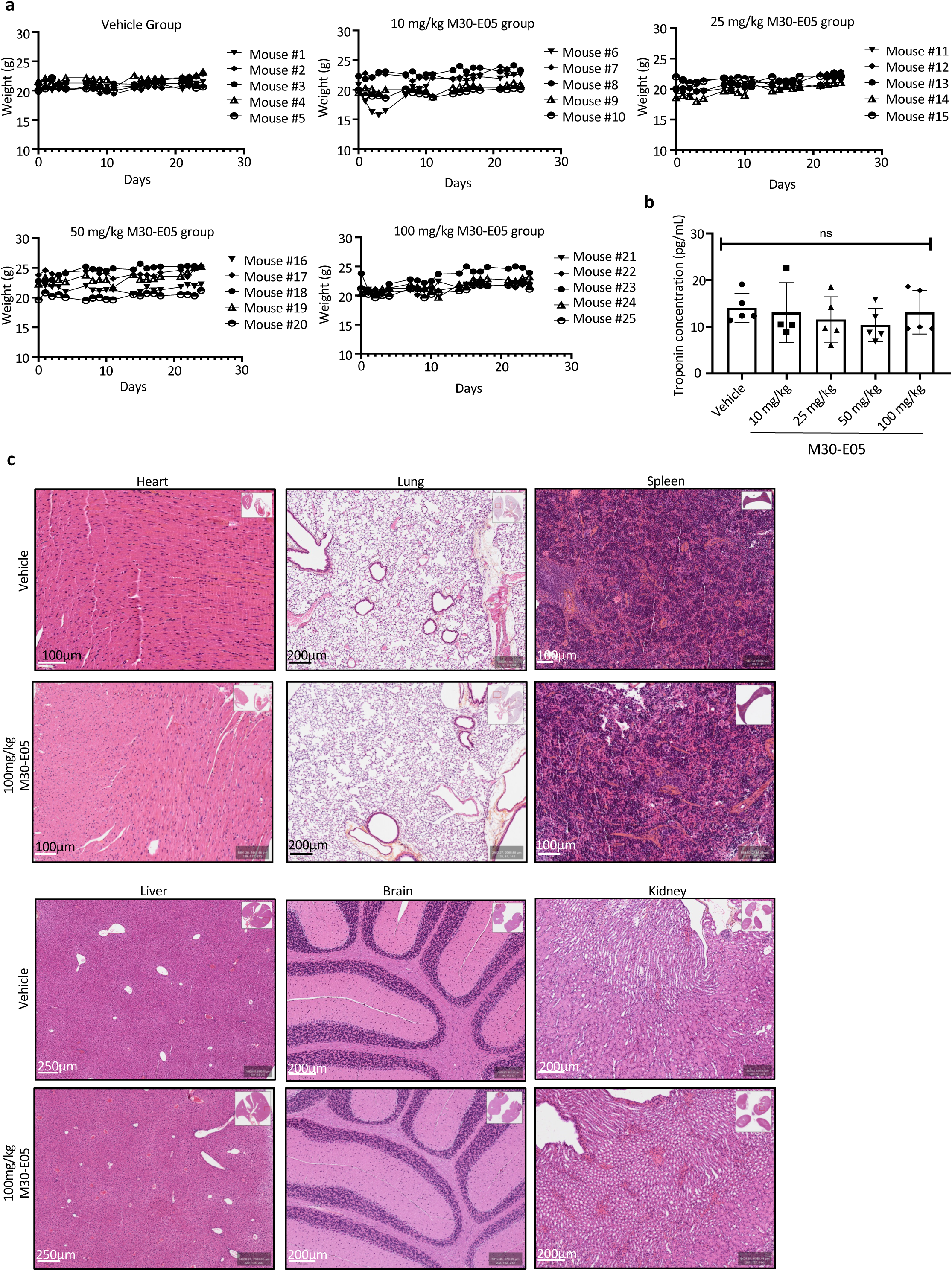
M30-E05 does not display any visible toxicity in non-engrafted NSG mice. **a,** Body weight curves of NSG mice orally administered with 0.5% methylcellulose (Vehicle), 10 mg/kg M30-E05, 25 mg/kg M30-E05, 50 mg/kg M30-E05, or 100 mg/kg M30-E05, five days per week for 4 weeks. Body weight of 5 mice per group was recorded on treatment days. Each curve represents an individual mouse. **b,** Serum cardiac troponin levels were quantified using a Mouse Cardiac Troponin I (TNNI3) ELISA Kit. Statistical analysis was performed using one-way ANOVA with Sidak’s multiple comparisons test. Significance levels are indicated as follows: ns, not significant; p < 0.0332 (*);p < 0.0021 (**); p < 0.0002 (***);p < 0.0001 (****). **c,** Hematoxylin/eosin (H&E) staining of heart, lung, spleen, liver, brain, and kidney tissues from NSG mice orally administered 0.5% methylcellulose (Vehicle) or 100 mg/kg M30-E05, five days per week for 4 weeks.

Prompted by the absence of M30-E05 toxicity, we next evaluated the anti-tumor efficacy of M30-E05 in the PDX GR-OS-18, which expressed the high level of AIF and CHCHD4, Fig. 5b). GR-OS-18 fragments were well subcutaneously established in NSG mice (80-150 mm^3^ tumor volume obtained 3 days after tumor transplantation). Mice were randomly assigned to M30-E05 treatment at 100 mg/kg, administered via gavage 5 days per week for 14 days (Fig. 7a). Vehicle-only treated mice served as controls. No effects on body weight were observed (Fig. 7b), M30-E05 caused a significant reduction of tumor growth by 41% after 2 weeks of treatment (Fig. 7c). Immunohistological staining revealed an M30-E05-induced decrease in Ki-67 expression, the mitotic index biomarker (Supplementary Fig. 5a) and a reduction of AIF protein expression, suggesting an on-target effect of the drug (Fig. 7d).

**Figure 7.**
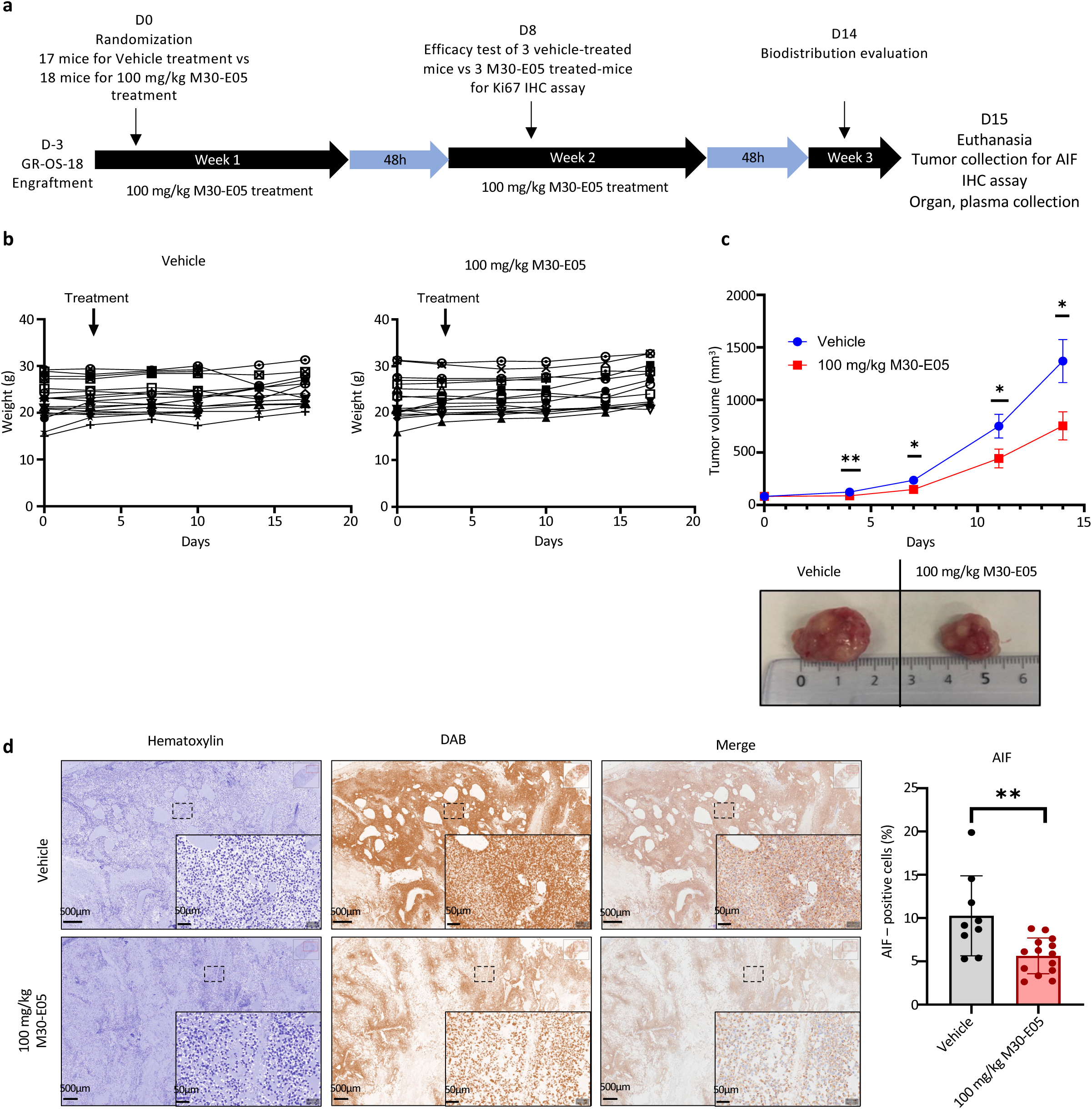
M30-E05 inhibits tumor growth in GR-OS-18 engrafted NSG mice. **a)** Experimental design. Mice were engrafted subcutaneously with GR-OS-18 PDX cells. Three days later, mice were orally administrated with 0.5% methylcellulose (Vehicle) or 100mg/kg M30-E05 five days per week for 2 weeks. The timing of sample collection for efficacy study, biodistribution analysis and sacrifice is indicated. IHC, immunochemistry. **b)** Body weight curves of GR-OS-18 engrafted NSG mice administered 0.5% methylcellulose (Vehicle) or 100 mg/kg M30-E05, five days per week for 2 weeks. Body weight (g) was recorded twice per week. Each curve represents an individual mouse. Vehicle group, n= 17 mice and 100 mg/kg M30-E05 treated group, n = 18 mice. **c)** Tumor volume curves of engrafted mice treated with 0.5% methylcellulose (Vehicle) or 100 mg/kg M30-E05. Vehicle group, n= 17 mice and 100 mg/kg M30-E05 treated group, n = 18 mice. Representative dissected tumors at the end of the experiment derived from vehicle and 100 mg/kg M30-E05-treated mice were shown. Data were shown as mean ± SD and statistical significance was determined by Mann-Whitney test for each time point. Significance levels are indicated as follows: ns, not significant; p < 0.0332 (*);p < 0.0021 (**); p < 0.0002 (***);p < 0.0001 (****). **d)** Immunochemistry (IHC) analysis of tumor sections for AIF expression from GR-OS-18 engrafted NSG mice administered 0.5% methylcellulose (Vehicle) or 100 mg/kg M30-E05, five days per week for 2 weeks. Vehicle group, n=9 mice and 100 mg/kg M30-E05 treated group, n = 1’ mice. Data were shown as mean ± SD and statistical significance was determined by Mann-Whitney test. Significance levels are indicated as follows: ns, not significant; p < 0.0332 (*);p < 0.0021 (**); p < 0.0002 (***);p < 0.0001 (****).

Mass spectrometric detection of M30-E05 was performed 2h and 4h following oral administration to mice. The compound was detected only in the bones and not in the spleen, liver, kidney, lung, plasma, heart and brain (Supplementary Fig. 5b). These findings align with the known pharmacokinetics of flavonoid, M30-E05 being rapidly metabolized following administration and can attain specific organs such as the bones.

## Discussion

Metabolic reprogramming is a hallmark of many cancers and targeting this vital process may hold promise to design new treatments for resistant cancer patients. With the aim to discover small molecules capable of modulating metabolic flexibility of malignant cells, we identified a novel engineered flavonoid, M30-E05. *In vitro*, this agent kills cells through the disruption of the AIF/CHCHD4 complex and apoptosis induction. Remarkably, M30-E05 is orally bioavailable and mediates significant tumor growth reduction in mice bearing PDX from a refractory osteosarcoma patient.

Flavonoids form a class of natural compounds well-known for their potent anti-inflammatory and neuroprotective properties that also bear potential as anticancer agents. For recent examples, the flavonoids *Cirsiliol*^32^ and *Baicalin*^33^ induce apoptosis in osteosarcoma cells and promote osteogenic differentiation *in vitro*, respectively. Using similar or lower doses than these compounds, we show that M30-E05 can be orally administered and then can redistribute into internal organs such as bones, though fails to accumulate into the heart and the brain. This characteristic represents a significant advantage, by minimizing the risk of cerebral and cardiac toxicity, which is a common concern for drugs that inhibit oxidative phosphorylation. Of note, a recent selective complex I inhibitor failed in two Phase I trials for advanced solid tumors and acute myeloid leukemia^34^.

We found that M30-E05 preferentially binds to the NAD-binding pocket of both the AIF monomer and dimer, suggesting that the molecule may competitively inhibit NAD binding, thereby preventing AIF dimerization and CHCHD4 interaction. Therefore, our compound exhibits a distinct mechanism compared to the recently identified 4-aminoquinolines, which stabilize AIF dimer and stimulate its interaction with CHCHD4^35^. Among the hits identified, M30-E05, which exhibited the most promising properties *in vitro* and *in cellulo*, was selected over 15 hits for further preclinical evaluation. Our study demonstrates that the biological mechanism of action of M30-E05 involves several key steps. First, M30-E05 disrupts the AIF/CHCHD4 complex, leading to a reduction in the expression levels of both AIF and CHCHD4, as well as that of their substrates COX17 and MICU1, the mitochondrial import of which depends on AIF/CHCHD4. These changes point to a disruption of mitochondrial protein import that then impacts the assembly of respiratory chain complexes. Accordingly, cells treated with M30-E05 exhibited alterations in mitochondrial network structure and function. This was evidenced by a significant reduction in mitochondrial oxygen consumption and ATP level, accompanied by a strong shift in energy substrate utilization from glucose to lipids and amino acids. Given these major changes, it is not surprising that M30-E05 compromised cell viability, thus inducing apoptosis.

Leveraging access to RNA-seq data from patients at different stages of tumor development (diagnosis, therapy resistance and relapse) as well as PDX models and both parental and resistant cell lines, we focused on osteosarcoma to preclinically evaluate the therapeutic potential of our drug candidate, M30-E05. Among pediatric cancers, osteosarcoma (predominantly affects children and young adults^36^) is characterized by high malignancy and a strong propensity for early metastasis, primarily to the lungs^37^ ^38^. Current treatment options are limited to chemotherapy and surgical resection^39^. However, despite significant progress in pediatric oncology and advancements in precision medicine, no improvement in survival rates of patients with osteosarcoma has been achieved over the past four decades. Consequently, a substantial proportion of patients with osteosarcoma remain uncured due to chemotherapy resistance or relapse^40^. Importantly, almost no mutations of *AIFM1* were found in any osteosarcoma patients, suggesting that AIF would not exhibit changes in amino acids sequence that could modify the accessibility of the protein to a small molecule and the bioactivity of the compound. Interestingly, our results suggest that M30-E05 is not a drug efflux pump substrate and overcomes drug resistance via the specific targeting of AIF/CHCHD4 complex.

RNA-seq analysis of PDX models and osteosarcoma cell lines revealed that mitochondrial function and energetic metabolism are reprogrammed and might be linked to the degree of chemoresistance, metastasis and tumor aggressiveness^41^ ^42^. Accordingly, the morphology of the mitochondrial network, mitochondrial respiration and expression of AIF and CHCHD4 and some of their substrates also changed in these models. Importantly, M30-E05 demonstrated the ability to kill a variety of osteosarcoma cancer cell lines and PDX models, irrespective of their multi-chemoresistant status, with IC_50_ values ∼20 µM for most PDX models. M30-E05 effects varied with the grafting site in mice suggesting a potential role of the tumoral microenvironment that remains to be investigated. At comparable doses, M30-E05 also exhibited cytotoxic activity against other cancer cell lines, including ovarian, lung, and leukemia, while showing limited toxicity on HEK293, HK2 cells and primary human PBMCs. This suggests that M30-E05 holds potential for broader application in adult cancers.

*In vivo* effects of M30-E05 were evaluated in immunodeficient NSG mice. The compound was formulated in 0.5% methylcellulose and administered via oral gavage five days per week for 4 weeks. M30-E05 is chemically modified to have an improved bioavailability and consistent with its chemical properties, M30-E05 was rapidly metabolized, well tolerated and found to be non-toxic to major murine organs at doses up to 100 mg/kg. This would not be due to a failure of M30-E05 to bind murine AIF since the docking experiments confirmed that the compound could bind similarly murine and human AIF. At 100 mg/kg, M30-E05 induced a significant tumor growth inhibition of 41%. Interestingly, tumor growth inhibition was accompanied by a decrease in AIF expression, as demonstrated by immunohistochemistry (IHC). If confirmed, this could position AIF as a potential biomarker of treatment responses.

The efficacy range of M30-E05 is a critical parameter for its drug development. The compound showed cytotoxic activity in the range of 2.5 to 12 µM across osteosarcoma and adult cancer cell lines, which, while promising, may be considered relatively high from a clinical perspective. Among the tested cell lines, ovarian cancer and leukemia cells demonstrated the greatest sensitivity, consistent with their metabolic reprogramming profiles^43^ ^44^. To improve M30-E05 efficacy, several strategies could be pursued: (i) developing improved formulations (e.g., nanoparticles or antibody-drug conjugates), (ii) designing more potent analogs, (iii) identifying more sensitive cancer types or subtypes for targeted development, or (iv) combining with other treatments such as radiotherapy and/or immunotherapy.

In conclusion, M30-E05, a novel chemically modified flavonoid, acts through an original biological mechanism targeting cancer-specific metabolic reprogramming via AIF/CHCHD4 inhibition. This compound might become a valuable therapeutic option for refractory and relapsed cancer patients, either as a monotherapy or in combination treatments.

## Acknowledgments

CB research was supported by grants from the French National Cancer Institute “INCa 2017-1-PL BIO-08” and “2021 - 167/ INCA_16344” and the Société française de lutte contre les cancers et les leucémies de l’enfant et de l’adolescent (SFCE), grant number ECS 20, PHC PESSOA, N°49163TL and the Groupement des Entreprises Francaises dans la lutte contre le Cancer [2023-2024]. BG work including the MAPPYACTS trial was supported by grants from INCa through the PHRC “INCa-DGOS_8519” MERRI, Fondation ARC, Association Imagine for Margo, Fédération Enfants et Santé, SFCE, and Dell. The MAPPYACTS PDX establishment was supported by grants from Fondation Gustave Roussy; Fédération Enfants Cancers et Santé, Société Française de lutte contre les Cancers et les leucémies de l’Enfant et l’adolescent (SFCE), Association AREMIG and Thibault BRIET; Parrainage médecin-chercheur of Gustave Roussy. AM and BG were supported by the “Parrainage médecin-chercheur” of Gustave Roussy. The work has been produced in collaboration with the BoOST-dataS consortium, coordinated by Gustave Roussy and including the Toulouse University Hospital, the UMR1238 unit of the University of Nantes, the Léon Bérard centre, the Curie Institute, Unicancer, the Strasbourg University Hospital & University, the CRBM of the University of Montpellier and the Oscar Lambret centre, and with the support of the French National Cancer Institute. GK is supported by the Ligue contre le Cancer (équipe labellisée); Agence National de la Recherche (ANR-22-CE14-0066 VIVORUSH, ANR-23-CE44-0030 COPPERMAC, ANR-23-R4HC-0006 Ener-LIGHT); Association pour la recherche sur le cancer (ARC); Cancéropôle Ile-de-France; Fondation pour la Recherche Médicale (FRM); a donation by Elior; European Joint Programme on Rare Diseases (EJPRD) Wilsonmed; European Research Council Advanced Investigator Award (ERC-2021-ADG, Grant No. 101052444; project acronym: ICD-Cancer, project title: Immunogenic cell death (ICD) in the cancer-immune dialogue); The ERA4 Health Cardinoff Grant Ener-LIGHT; European Union Horizon 2020 research and innovation programmes Oncobiome (grant agreement number: 825410, Project Acronym: ONCOBIOME, Project title: Gut OncoMicrobiome Signatures [GOMS] associated with cancer incidence, prognosis and prediction of treatment response, Prevalung (grant agreement number 101095604, Project Acronym: PREVALUNG EU, project title: Biomarkers affecting the transition from cardiovascular disease to lung cancer: towards stratified interception), Neutrocure (grant agreement number 861878 : Project Acronym: Neutrocure ; project title: Development of “smart” amplifiers of reactive oxygen species specific to aberrant polymorphonuclear neutrophils for treatment of inflammatory and autoimmune diseases, cancer and myeloablation); National support managed by the Agence Nationale de la Recherche under the France 2030 programme (reference number 21-ESRE-0028, ESR/Equipex+ Onco-Pheno-Screen); Hevolution Network on Senescence in Aging (reference HF-E Einstein Network); Institut National du Cancer (INCa); Institut Universitaire de France; LabEx Immuno-Oncology ANR-18-IDEX-0001; a Cancer Research ASPIRE Award from the Mark Foundation; PAIR-Obésité INCa_1873, the RHUs Immunolife and LUCA-pi (ANR-21-RHUS-0017 and ANR-23-RHUS-0010, both dedicated to France Relance 2030); Seerave Foundation; SIRIC Cancer Research and Personalized Medicine (CARPEM, SIRIC CARPEM INCa-DGOS-Inserm-ITMO Cancer_18006 supported by Institut National du Cancer, Ministère des Solidarités et de la Santé and INSERM). Tudor Manolu (PFIC Platform) and Sylvie Souquère (EM platform) from UMS AMMICA, Gustave Roussy, Villejuif, Manon Cruette (INSERM U1180, Univ Paris-Saclay, Châtenay Malabry), Delphine Courilleau (CIBLOT platform Univ Paris-Saclay, Gif Sur Yvette) are acknowledged for their technical help.

## Conflicts of interest

GK has been holding research contracts with Daiichi Sankyo, Eleor, Kaleido, Lytix Pharma, PharmaMar, Osasuna Therapeutics, Samsara Therapeutics, Sanofi, Sutro, Tollys, and Vascage. GK is on the Board of Directors of the Bristol Myers Squibb Foundation France. GK is a scientific co-founder of everImmune, Osasuna Therapeutics, Samsara Therapeutics and Therafast Bio. GK is in the scientific advisory boards of Centenara Labs (formerly Rejuveron Life Sciences), Hevolution, and Institut Servier. GK is the inventor of patents covering therapeutic targeting of aging, cancer, cystic fibrosis and metabolic disorders. Among these, patents were licensed to Bayer (WO2014020041-A1, WO2014020043-A1), Bristoll-Myers Squibb (WO2008057863-A1), Osasuna Therapeutics (WO2019057742A1), PharmaMar (WO2022049270A1 and WO2022048775-A1), Raptor Pharmaceuticals (EP2664326-A1), Samsara Therapeutics (GB202017553D0), and Therafast Bio (EP3684471A1). GK’wife, Laurence Zitvogel, has held research contracts with Glaxo Smyth Kline, Incyte, Lytix, Kaleido, Innovate Pharma, Daiichi Sankyo, Pilege, Merus, Transgene, 9 m, Tusk and Roche, was on the on the Board of Directors of Transgene, is a cofounder of everImmune, and holds patents covering the treatment of cancer and the therapeutic manipulation of the microbiota. Among these, patents were licensed to everImmune (US20210346438A1, US20200360449A1) and Transgene (US20200376052A1). GK’s brother, Romano Kroemer, was an employee of Sanofi and now consults for Boehringer-Ingelheim. The funders had no role in the design of the study; in the writing of the manuscript, or in the decision to publish the results.

Nathalie Gaspar has got consultant activity for EISAI and IPSEN, and participate to advisory board for ABBVIE, MERCKS, Y-MABS, Daiichi Sankyo/AstraZeneca.

Figures were created with BioRender.com

**Supplemental Figure 1. AIF and CHCHD4 expression levels in patients, PDX and cell lines.**

**a, b,** *AIFM1* and *CHCHD4* mRNA expression was determined in patients at diagnosis (OS2006) vs relapse (MAPPYACTS) and **c,** in 8 PDX xenografted subcutaneously (sc) and paratibial (Ortho) in tpm (transcript per million). Data were shown as mean ± SD and statistical significance was determined by Mann-Whitney test for each time point. Significance levels are indicated as follows: ns, not significant; *p < 0.0332; **p < 0.0021; ***p < 0.0002; **p < 0.0001.

**c,** Expression of AIF and CHCHD4 in HOS cells, HOS-R/DOXO and HO-R/MTX cells were analyzed by Western-blot with VDAC as loading control and protein expression was quantified in ratio to VDAC. A representative western-blot is presented while the experiment was repeated 3 times. VDAC was used as a loading control.

**Supplementary Figure 2. Mitochondrial network metabolic activity is altered by drug-induced resistance in HOS cells.**

**a,** Representative confocal images of HOS, HOS-R/DOXO and HOS-R/MTX’s mitochondrial network. Mitochondria are labelled with TOM20 antibody and nuclei were stained with DAPI. Four independent experiments were performed, and Z-stack projections of each condition were evaluated.

**b,** Representative transmission electron microscopy images of HOS, HOS-R/DOXO and HOS-R/MTX’s cells, showing differences in mitochondria shape and cristae density. Ratios between the length and width of mitochondria of each condition were measured (n>100). Data is represented as violon plots, and the median of each condition is shown by the horizontal dashed line. A ration equal to 1 indicates that mitochondria have a round shape. Statistical significance was assessed using ANOVA with Dunnett’s multiple comparison test correction. Significance levels are indicated as follows: p < 0.01 (**), and p < 0.0001 (****) *vs* HOS.

**c,** Mitochondrial membrane potential assessed by flow cytometry using 100nM TMRM. fluorescent probe labelling for 20 min. At least 4 independent experiments were performed (n≥4). Data represent the mean±SD. Statistical significance was assessed using ANOVA with Dunnett’s multiple comparison test correction. Significance levels are indicated as follows: < 0.01 (**) *vs* HOS.

**d,** Seahorse XFe96 Mito Stress Test graph displaying the oxygen consumption rate (OCR; pmol/min/1.000 cells) over time in HOS, HOS-R/DOXO and HOS-R/MTX cells. The test was conducted using the following compounds: Oligomycin (2.5 µM), FCCP (1 µM), and Rotenone/Antimycin A (0.5 µM). Three independent experiments were performed.

**e,** ATP levels were semi-quantified over time in HOS, HOS-R/DOXO and HOS-R/MTX cells, using the ATPlite assay (n ≥4). Data represent the mean±SD. Statistical analysis was performed using ANOVA with Dunnett’s multiple comparison test correction. Significance levels are indicated as follows: p < 0.0001 (****) vs HOS.

**f, g,** Enrichment analysis of metabolites that differ significantly between HOS and HOS-R/DOXO **f,** and between HOS and HOS-R/MTX **g,** cell lines. The enrichment ratio represents the observed number of metabolites in a specific metabolic pathway divided by the expected number. Metabolic pathways are ranked by p-value, with the most significant pathways at the top.

**Supplementary Figure 3. Reponses to standard treatments of osteosarcoma and mitochondrial ultrastructure in secondary cultured PDX samples**

**a, b,** Cell viability of 6 different secondary cultured in vitro PDX samples treated with **a,** Cisplatin and **b,** Etoposide – osteosarcoma standard clinical treatments, measured using an LDH assay. Dose-response curves represent cell viability at increasing concentrations of M30-E05 after 72h of treatment. Individual replicates are shown (n > 2).

**c,** Representative transmission electron microscopy images of 6 different secondary cultured in vitro PDX samples in basal condition, showing different unique mitochondrial ultrastructure. m: mitochondrion

**Supplementary Figure 4. Immunofluorescence analysis of AIF, CHCHD4 and MICU1 expression in PDX tumor sections**

Immunofluorescence analysis of tumor sections for **a,** AIF; **b,** CHCHD4; MICU1 **c,** expression from all 6 PDX models engrafted NSG mice. Blue fluorescence was stained to locate nucleus by Hoechst 33342, orange fluorescence was stained to mitochondria by MTC antibody, green fluorescence was stained to our target (AIF, CHCHD4 or MICU1) by their specific antibody. Digital analysis was performed using QuPath software.

**Supplementary Figure 5. Ki67 expression in GR-OS-18 tumor sections following treatment with M30-E05 and Biodistribution of M30-E05 in several organs**

**a,** IHC analysis of tumor sections for Ki67 expression from 6 GR-OS-18 engrafted NSG mice administered 0.5% methylcellulose (Vehicle) (n =3) or 100 mg/kg M30-E05 (n = 3), five days per week for 2 weeks. 3 Vehicle-treated mice and 3 100 mg/kg M30-E05 treated-mice were euthanized at day 8 of treatment. Digital analysis was performed using QuPath software. Data were shown as mean ± SD and statistical significance was determined by Mann-Whitney test.

**b,** Biodistribution of M30-E05 in mouse brain, lung, heart, spleen, kidney, bone and liver after oral administration of 100 mg/kg M30-E05 at 2h and 4h post-administration. The organs were sampled for biodistribution analysis with liquid chromatography-tandem mass spectrometry (LC-MS/MS).

## References

1. Warburg, O., Wind, F. & Negelein, E. THE METABOLISM OF TUMORS IN THE BODY. J Gen Physiol 8, 519–530 (1927).

2. Hanahan, D. & Weinberg, R. A. Hallmarks of cancer: the next generation. Cell 144, 646– 674 (2011).

3. Faubert, B., Solmonson, A. & DeBerardinis, R. J. Metabolic reprogramming and cancer progression. Science 368, eaaw5473 (2020).

4. Hanahan, D. Hallmarks of Cancer: New Dimensions. Cancer Discov 12, 31–46 (2022).

5. Lai, H. T. et al. Insight into the interplay between mitochondria-regulated cell death and energetic metabolism in osteosarcoma. Front Cell Dev Biol 10, 948097 (2022).

6. Reinhardt, C. et al. AIF meets the CHCHD4/Mia40-dependent mitochondrial import pathway. Biochim Biophys Acta Mol Basis Dis 1866, 165746 (2020).

7. Benihoud, K. & Brenner, C. Profiling the Energy Metabolism at the Cell Subpopulation Level. J Cancer Immunol Volume 3, 147–150 (2021).

8. Rath, S. et al. MitoCarta3.0: an updated mitochondrial proteome now with sub-organelle localization and pathway annotations. Nucleic Acids Research 49, D1541–D1547 (2021).

9. Meyer, K. et al. Loss of apoptosis-inducing factor critically affects MIA40 function. Cell Death Dis 6, e1814 (2015).

10. Petrungaro, C. et al. The Ca(2+)-Dependent Release of the Mia40-Induced MICU1-MICU2 Dimer from MCU Regulates Mitochondrial Ca(2+) Uptake. Cell Metab 22, 721– 733 (2015).

11. Wang, T. et al. C9orf72 regulates energy homeostasis by stabilizing mitochondrial complex I assembly. Cell Metab 33, 531–546.e9 (2021).

12. Salscheider, S. L. et al. AIFM1 is a component of the mitochondrial disulfide relay that drives complex I assembly through efficient import of NDUFS5. EMBO J 41, e110784 (2022).

13. Nedara, K. et al. Relevance of the TRIAP1/p53 axis in colon cancer cell proliferation and adaptation to glutamine deprivation. Front Oncol 12, 958155 (2022).

14. Wischhof, L., Scifo, E., Ehninger, D. & Bano, D. AIFM1 beyond cell death: An overview of this OXPHOS-inducing factor in mitochondrial diseases. EBioMedicine 83, 104231 (2022).

15. Rao, S. et al. AIF-regulated oxidative phosphorylation supports lung cancer development. Cell Res 29, 579–591 (2019).

16. Al-Habib, H. & Ashcroft, M. CHCHD4 (MIA40) and the mitochondrial disulfide relay system. Biochem Soc Trans 49, 17–27 (2021).

17. Brenner, C. & Modjtahedi, N. Method for Identifying Compounds Useful for the Treatment of Cancer. (2021).

18. Hangen, E. et al. Interaction between AIF and CHCHD4 Regulates Respiratory Chain Biogenesis. Mol Cell 58, 1001–1014 (2015).

19. Marques da Costa, M. E., et al. In-Vitro and In-Vivo Establishment and Characterization of Bioluminescent Orthotopic Chemotherapy-Resistant Human Osteosarcoma Models in NSG Mice. Cancers (Basel) 11, 997 (2019).

20. Marques Da Costa, M. E., et al. A biobank of pediatric patient-derived-xenograft models in cancer precision medicine trial MAPPYACTS for relapsed and refractory tumors. Commun Biol 6, 949 (2023).

21. Schindelin, J., et al. Fiji: an open-source platform for biological-image analysis. Nat Methods 9, 676–682 (2012).

22. Bolte, S. & Cordelières, F. P. A guided tour into subcellular colocalization analysis in light microscopy. J Microsc 224, 213–232 (2006).

23. Ye, H. et al. DNA binding is required for the apoptogenic action of apoptosis inducing factor. Nat Struct Biol 9, 680–684 (2002).

24. Waterhouse, A. et al. SWISS-MODEL: homology modelling of protein structures and complexes. Nucleic Acids Res 46, W296–W303 (2018).

25. Abraham, M. J. et al. GROMACS: High performance molecular simulations through multi-level parallelism from laptops to supercomputers. SoftwareX 1–2, 19–25 (2015).

26. Lindorff-Larsen, K. et al. Improved side-chain torsion potentials for the Amber ff99SB protein force field. Proteins 78, 1950–1958 (2010).

27. Jorgensen, W. L., Chandrasekhar, J., Madura, J. D., Impey, R. W. & Klein, M. L. Comparison of simple potential functions for simulating liquid water. The Journal of Chemical Physics 79, 926–935 (1983).

28. Trott, O. & Olson, A. J. AutoDock Vina: improving the speed and accuracy of docking with a new scoring function, efficient optimization, and multithreading. J Comput Chem 31, 455–461 (2010).

29. Maté, M. J. et al. The crystal structure of the mouse apoptosis-inducing factor AIF. Nat Struct Mol Biol 9, 442–446 (2002).

30. Berlanga, P. et al. The European MAPPYACTS Trial: Precision Medicine Program in Pediatric and Adolescent Patients with Recurrent Malignancies. Cancer Discov 12, 1266– 1281 (2022).

31. Marchais, A. et al. Immune Infiltrate and Tumor Microenvironment Transcriptional Programs Stratify Pediatric Osteosarcoma into Prognostic Groups at Diagnosis. Cancer Res 82, 974–985 (2022).

32. Luo, M. et al. Cirsiliol induces autophagy and mitochondrial apoptosis through the AKT/FOXO1 axis and influences methotrexate resistance in osteosarcoma. J Transl Med 21, 907 (2023).

33. Zhao, Z.-F., Cheng, X.-F., Yu, M., Shi, W.-F. & Zhang, T. Baicalin Plays an Anti-Osteosarcoma Role in Vitro and Promotes Osteogenic Differentiation by Inhibiting NF-κB Signaling. Discov Med 36, 1648–1656 (2024).

34. Yap, T. A. et al. Complex I inhibitor of oxidative phosphorylation in advanced solid tumors and acute myeloid leukemia: phase I trials. Nat Med 29, 115–126 (2023).

35. Brosey, C. A. et al. Chemical screening by time-resolved X-ray scattering to discover allosteric probes. Nat Chem Biol 20, 1199–1209 (2024).

36. Gill, J. & Gorlick, R. Advancing therapy for osteosarcoma. Nat Rev Clin Oncol 18, 609–624 (2021).

37. Mirabello, L., Troisi, R. J. & Savage, S. A. Osteosarcoma incidence and survival rates from 1973 to 2004: data from the Surveillance, Epidemiology, and End Results Program. Cancer 115, 1531–1543 (2009).

38. Bielack, S. S. et al. Prognostic factors in high-grade osteosarcoma of the extremities or trunk: an analysis of 1,702 patients treated on neoadjuvant cooperative osteosarcoma study group protocols. J Clin Oncol 20, 776–790 (2002).

39. Gaspar, N. et al. Results of methotrexate-etoposide-ifosfamide based regimen (M-EI) in osteosarcoma patients included in the French OS2006/sarcome-09 study. Eur J Cancer 88, 57–66 (2018).

40. Casali, P. G. et al. Bone sarcomas: ESMO-PaedCan-EURACAN Clinical Practice Guidelines for diagnosis, treatment and follow-up. Ann Oncol 29, iv79–iv95 (2018).

41. Wei, Q., Qian, Y., Yu, J. & Wong, C. C. Metabolic rewiring in the promotion of cancer metastasis: mechanisms and therapeutic implications. Oncogene 39, 6139–6156 (2020).

42. Demicco, M., Liu, X.-Z., Leithner, K. & Fendt, S.-M. Metabolic heterogeneity in cancer. Nat Metab 6, 18–38 (2024).

43. Wang, M., Zhang, J. & Wu, Y. Tumor metabolism rewiring in epithelial ovarian cancer. J Ovarian Res 16, 108 (2023).

44. Feng, L., Zhang, P. Y., Gao, W., Yu, J. & Robson, S. C. Targeting chemoresistance and mitochondria-dependent metabolic reprogramming in acute myeloid leukemia. Front Oncol 13, 1244280 (2023).

