## Supplementary material for "Engineered flavonoid disrupts mitochondrial AIF/CHCHD4 complex for targeted cancer therapy": supp note

**Synthesis of M30-E05 : Protocols and Products Characterizations**

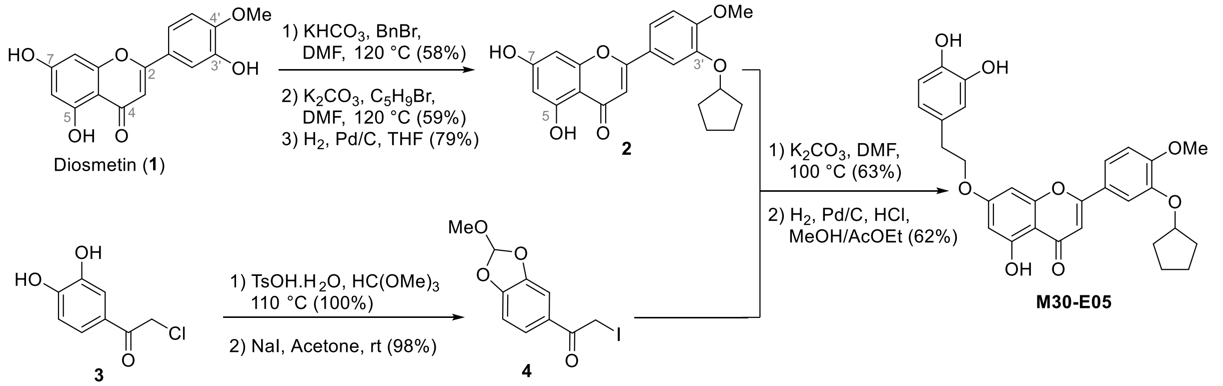


<u>3’-O-cyclopentyl-diosmetin **2**</u>

To a solution of diosmetin (21.0 g, 70 mmol, 1 equiv) in DMF (200 mL) was added $KHCO_3$ (7g, 70 mmol, 1 equiv). The reaction mixture was heated at 120 °C and benzyl bromide (12.5 mL, 105 mmol, 1.5 equiv) was added and the reaction mixture was heated for 150 min at that temperature. TLC monitoring indicated that starting material was remaining, but the reaction was stopped to keep selectivity. After cooling the reaction to rt, 1:1 AcOEt: Toluene mixture (200 mL) was added and the reaction mixture was filtered over celite, washing with the celite pad with 1:1 AcOEt: Toluene mixture. After concentration to dryness on a rotavapor, hot EtOH (200 mL) was added and the reaction mixture was let crystallizing over week-end at rt. The crystals were recovered by suction, washed with EtOH, $Et_2O$ and pentane. After drying, 7-O-benzyl-diosmetin was recovered as yellow crystals (16.0 g, 58%), contaminated with about 8% of starting material.

**Mp :** 204-206 °C. **$^1$H NMR** (300 MHz, THF-$d_8$) δ 12.97 (s, 1H), 8.28 (s, 1H), 7.52 – 7.42 (m, 4H), 7.42 – 7.25 (m, 4H), 7.02 (d, *J* = 8.3 Hz, 1H), 6.66 (s, 1H), 6.40 (d, *J* = 2.0 Hz, 1H), 5.19 (s, 2H), 3.92 (s, 3H). **$^{13}$C NMR** (75 MHz, THF) δ 183.2, 165.7, 165.1, 163.6, 158.8, 152.0, 148.4, 137.7, 129.4, 129.3, 128.9, 128.5, 125.2, 119.3, 114.1, 112.1, 105.1, 99.5, 93.9, 71.2, 56.4. **IR** ($cm^{-1}$) : 3346, 1655, 1602, 1495, 1431, 1356, 1250, 1157, 1115, 1022, 829, 763. **HRMS-ESI** (*m /z*) : calcd for $[M+H]^+$ $C_{23}H_{19}O_6$, 390.1182 ; Found : 390.1179.

To a solution of 7-O-benzyl-diosmetin (16.0 g, 41 mmol) in DMF (150 mL) was added $K_2CO_3$ (7.4 g, 53.3 mmol, 1.3 equiv). The reaction mixture was heated at 120 °C, cyclopentyl bromide (6.2 mL, 61.5 mmol, 1.5 equiv.) was added and the reaction mixture was stirred at that temperature for 5 h, until complete conversion by TLC monitoring. The reaction mixture was cooled down to rt, and was quenched by water (200 mL) and diluted with AcOEt (400 mL). The organic layer was washed 3 times with 1N HCl, and then with brine. The organic layer was dried over $MgSO_4$, filtered, concentrated and the residue was recrystallized with hot MeOH to give 3’-O-cyclopentyl-7-O-benzyl-diosmetin as yellow/brown crystals (11.0 g, 59%).

**Mp** : 200-202 °C **$^1$H NMR** (300 MHz, Chloroform-*d*) δ 12.82 (s, 1H), 7.48 (dd, *J* = 8.6, 2.2 Hz, 1H), 7.45 – 7.35 (m, 5H), 7.33 (d, *J* = 2.3 Hz, 1H), 6.95 (d, *J* = 8.6 Hz, 1H), 6.60 – 6.51 (m, 2H), 6.44 (d, *J* = 2.3 Hz, 1H), 5.13 (s, 2H), 4.85 (tt, *J* = 6.3, 3.3 Hz, 1H), 3.92 (s, 3H), 2.07 – 1.77 (m, 6H), 1.74 – 1.55 (m, 2H). **$^{13}$C NMR** (75 MHz, $CDCl_3$) δ 182.5, 164.6, 164.3, 162.3, 157.7, 153.5, 148.0, 135.9, 128.9, 128.5, 127.6, 123.7, 120.0, 112.4, 111.7, 105.8, 104.6, 98.8, 93.7, 80.9, 70.6, 56.2, 32.9, 24.2. **IR** ($cm^{-1}$) : 2951, 1655, 11585, 1497, 1354, 1252, 1157, 1020, 831, 766, 704, 627. **HRMS-ESI** (*m /z*) : calcd for $[M+H]^+$ $C_{28}H_{27}O_6$, 459.1808 ; Found : 459.1805.

To a solution of 3’-O-cyclopentyl-7-O-benzyl-diosmetin (11.0 g, 24.0 mmol, 1 equiv) in THF (200 mL) in a Parr reactor was added Pd/C 10% (1.0 g). The reaction mixture was stirred under $H_2$ atmosphere (10 bars) for 24 h under vigorous stirring at rt. The reaction mixture was then filtered on celite and concentrated. The residue was triturated with hot DCM (50 mL) and was let cooling in the freezer. The solid was then recovered by suction, washed with cold DCM, then with $Et_2O$. The solid was then dried under vacuum and pure 3’-O-cyclopentyl-diosmetin (**2**) was recovered as a light brown powder (7.0 g, 79 %).

**Mp** : 207-209 °C. **$^1$H NMR** (300 MHz, DMSO-$d_6$) δ 12.92 (s, 1H), 10.80 (brs, 1H), 7.65 (dd, *J* = 8.6, 2.2 Hz, 1H), 7.53 (d, *J* = 2.2 Hz, 1H), 7.11 (d, *J* = 8.6 Hz, 1H), 6.91 (s, 1H), 6.51 (d, *J* = 2.2 Hz, 1H), 6.20 (d, *J* = 2.2 Hz, 1H), 4.98 (tt, *J* = 5.8, 2.6 Hz, 1H), 3.84 (s, 3H), 2.03 – 1.84 (m, 2H), 1.81 – 1.65 (m, 4H), 1.65 – 1.49 (m, 2H).**$^{13}$C NMR** (75 MHz, DMSO) δ 181.7, 164.2, 163.3, 161.4, 157.3, 153.0, 147.2, 122.9, 119.9, 112.2, 112.1, 103.7, 98.8, 94.0, 79.8, 55.7, 32.1, 23.6. **IR** ($cm^{-1}$) : 2932, 1651, 1557, 1495, 1435, 1367, 1331, 1250, 1169, 1036, 835, 725, 640. **HRMS-ESI** (*m /z*) : calcd for $[M+H]^+$ $C_{21}H_{21}O_6$, 369.1338 ; Found : 369.1334.

<u>2-iodo-1-(2-methoxybenzo[d][1,3]dioxol-5-yl)ethan-1-one **4**</u>

To a solution of chloromethyl-3,4-dihydroxyphenyl) ketone (10.0 g, 53.8 mmol) in trimethyl orthoformate (100 mL) was added PTSA monohydrate (500 mg) and the mixture was refluxed 2 days. The reaction mixture was the cooled down to rt, diluted with $Et_2O$ (200 mL) washed with $K_2CO_3$ 10% solution (3x100 mL), dried over $MgSO_4$, filtered, and concentrated to afford pure orthoester-protected catechol (16.8 g, 100%).

**Mp** : 140-142 °C. **$^1$H NMR** (300 MHz, Chloroform-*d*) *δ* 7.59 (dd, *J* = 8.2, 1.8 Hz, 1H), 7.49 (d, *J* = 1.8 Hz, 1H), 6.93 (d, *J* = 8.1 Hz, 1H), 6.92 (s, 1H), 4.61 (s, 2H), 3.41 (s, 3H). **$^{13}$C NMR** (75 MHz, Chloroform-*d*) *δ* 189.5, 150.9, 147.0, 129.1, 124.9, 120.2, 108.2, 108.0, 50.4, 45.7. **HRMS-ESI** (*m /z*) : calcd for $[M+H]^+$ $C_{10}H_{10}ClO_4$, 229.0268 ; Found : 229.0258. **IR** ($cm^{-1}$) : 3313, 1687, 1600, 1497, 1445, 1337, 1250, 1198, 1107, 989, 928, 806, 746.

To a solution of previous protected catechol (12.8 g, 53.8 mmol) in acetone (100 mL), was added NaI (12.1 g, 80.7 mmol, 1.5 equiv) and the reaction mixture was stirred at rt for 3 h. The reaction mixture was quenched with water and diluted with ether. The aqueous layer was extracted twice with $Et_2O$, and the combined organic layers were washed twice with 10% $NaHSO_3$ solution, dried over $MgSO_4$, filtered, concentrated, and the residue was obtained as a light brown oil, which crystallizes slowly on standing (16.87 g, 98%).

**Mp** :  73-75 °C. **$^1$H NMR** (300 MHz, Chloroform-*d*) δ 7.63 (dd, *J* = 8.2, 1.8 Hz, 1H), 7.51 (d, *J* = 1.8 Hz, 1H), 6.93 (s, 1H), 6.92 (d, *J* = 8.1 Hz, 1H), 4.29 (s, 2H), 3.42 (s, 3H). **$^{13}$C NMR** (75 MHz, Chloroform-*d*) *δ* 191.3, 150.8, 146.9, 128.3, 125.5, 120.2, 108.6, 108.0, 50.4, 1.5. **IR** ($cm^{-1}$) : 1660, 1599, 1491, 1435, 1392, 1263, 1196, 1144, 1022, 910, 879, 837, 769. **HRMS-ESI** (*m /z*) : calcd for $[M+H]^+$ $C_{10}H_{10}IO_4$, 320.9624 ; Found : 320.9614.

<u>M30-E05</u>

To a solution of 3’-O-cyclopentyl-diosmetin (**2**) (7.0 g, 19.0 mmol) in DMF (90 mL) was added $K_2CO_3$ (2.6 g, 19.0 mmol, 1 equiv). The reaction mixture was heated at 100 °C, iodide **4** (6.1 g, 19.0 mmol) in DMF (10 mL) was then added. The reaction was stirred 2 h at 100 °C, TLC monitoring showing the consumption of both starting materials. The reaction mixture was thus cooled down to rt, quenched with water (200 mL), extracted with DCM (3x 150 mL), and the combined organic layers were washed with brine (3x150 mL). The organic phase was dried over $MgSO_4$, filtered, concentrated and the residue was purified grossly on a silica gel chromatography (DCM: AcOEt, 90:10). The orange residue was then triturated with boiling MeOH (100 mL), and the mixture was cooled down to rt. The beige solid was filtered on a sintered glass, washed with a small amount of cold MeOH, and dried under vacuum to afford 7-O-(2-(2-methoxybenzo[d][1,3]dioxol-5-yl)-3'-O-cyclopentyl-diosmetin as a yellowish powder (6.72 g, 63%).

**Mp** : 161-163 °C. **$^{1}$H NMR** (300 MHz, Chloroform-*d*) $\delta$ 12.83 (s, 1H), 7.65 (dd, $J$ = 8.2, 1.7 Hz, 1H), 7.54 (d, $J$ = 1.7 Hz,1H), 7.47 (dd, $J$ = 8.5, 2.2 Hz, 1H), 7.31 (d, $J$ = 2.2 Hz, 1H), 6.98 (d, $J$ = 8.1 Hz, 1H), 6.95 (s, 1H), 6.94 (d, $J$ = 8.5 Hz, 1H), 6.55 (s, 1H), 6.53 (d, $J$ = 2.3 Hz, 1H), 6.34 (d, $J$ = 2.2 Hz, 1H), 5.30 (s, 2H), 4.85 (tt, $J$ = 6.2, 2.9 Hz, 2H), 3.91 (s, 3H), 3.43 (s, 3H), 2.06 – 1.77 (m, 6H), 1.72 – 1.57 (m, 2H). **$^{13}$C NMR** (75 MHz, Chloroform-*d*) $\delta$ 191.4, 182.5, 164.48, 163.7, 162.4, 157.7, 153.5, 151.1, 148.0, 147.1, 129.0, 124.5, 123.6, 120.2, 120.1, 112.4, 111.7, 108.2, 107.9, 106.2, 104.7, 98.4, 93.8, 80.9, 70.6, 56.2, 50.4, 32.9, 24.2. **IR** (cm$^{-1}$) : 2955, 1693, 1654, 1597, 1497, 1437, 1354, 1253, 1177, 1093, 904, 768. **HRMS-ESI** (*m /z*) : calcd for $[M+H]^+$ $C_{31}H_{29}O_{10}$, 561.1761 ; Found : 561.1747.

To a suspension of 7-O-(2-(2-methoxybenzo[d][1,3]dioxol-5-yl)-3'-O-cyclopentyl-diosmetin (3.0 g, 5.35 mmol, 1 equiv) in AcOEt:THF:MeOH mixture (30:40:30 mL) in a Parr reactor was added $Pd(OH)_2$ 20% on carbon (500 mg) and HCl 35% (100 µL) was added. The whole mixture was stirred (1200 rpm) at rt under $H_2$ atmosphere (10 bars) for 15 h. The mixture was then filtered on Celite, and concentrated on a rotavapor. The residue was triturated in with hot MeOH (20 mL), cooled down and $Et_2O$ was added (20 mL), continuing the trituration at rt. The beige powder was then isolated by filtration to afford M30-E05 (1.32 g) and the filtrate was concentrated and purified grossly on a silica gel chromatography (DCM:Acetone, 9:1). The purified product was triturated again in a 1:1 MeOH:$Et_2O$ mixture to give additional M30-E05 (350 mg). The combined fraction gives M30-E05 as a light beige powder (1.67 g, 62%).

**Mp** : 205-207 °C. **$^{1}$H NMR** (300 MHz, DMSO-*d*$_6$) δ 12.92 (s, 1H), 8.72(s, 1H), 8.67 (s, 1H), 7.67 (dd, $J$ = 8.7, 1.2 Hz, 1H), 7.54 (d, $J$ = 1.2 Hz, 1H), 7.11 (d, $J$ = 8.7 Hz, 1H), 6.96 (s, 1H), 6.78 (d, $J$ = 1.7 Hz, 1H), 6.78 (d, $J$ = 1.2 Hz, 1H), 6.67 (d, $J$ = 8.0 Hz, 1H), 6.56 (dd, $J$ = 8.0, 1.2 Hz, 1H), 6.34 (d, $J$ = 1.7 Hz, 1H), 5.00 (tt, $J$ = 6.2, 1.7 Hz, 1H), 4.23 (t, $J$ = 6.9 Hz, 2H), 3.84 (s, 3H), 2.88 (t, $J$ = 6.9 Hz, 2H), 2.04 – 1.87 (m, 2H), 1.86 – 1.66 (m, 4H), 1.66 – 1.49 (m, 2H).**$^{13}$C NMR** (75 MHz, DMSO-*d*$_6$) $\delta$ 182.0, 164.4, 163.6, 161.2, 157.3, 153.1, 147.2, 145.1, 143.8, 128.5, 122.7, 120.1, 119.6, 116.4, 115.5, 112.1, 112.0, 104.7, 103.9, 98.3, 93.2, 79.8, 69.5, 67.0, 55.8, 32.2, 23.7. **IR** (cm$^{-1}$) : 3402, 3161, 2953, 1654, 1600, 1499, 1438, 1251, 1165, 1024, 849, 808, 765. **HRMS-ESI** (*m /z*) : calcd for $[M+H]^+$ $C_{29}H_{29}O_8$, 505.1862 ; Found : 505.1849.

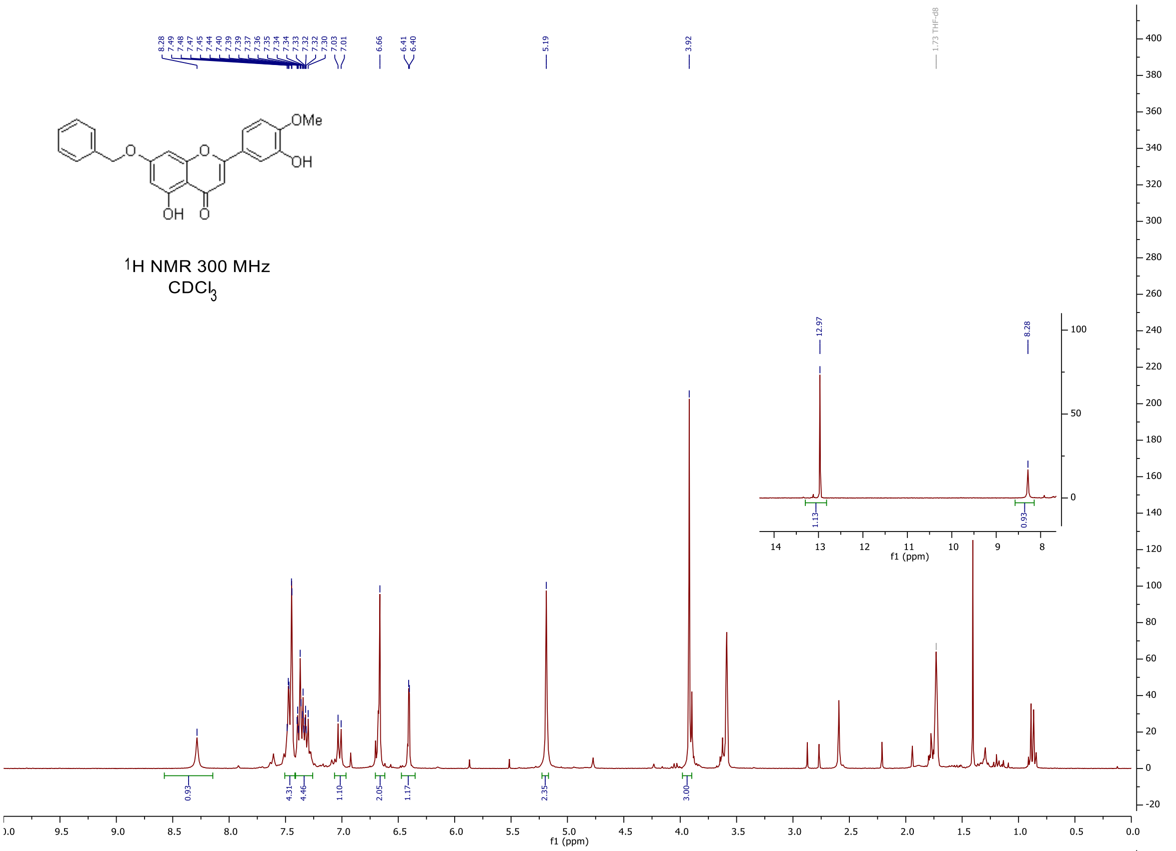
8.28 7.49 7.48 7.47 7.45 7.44 7.40 7.39 7.39 7.37 7.36 7.35 7.34 7.34 7.33 7.32 7.32 7.30 7.03 7.01
6.66
6.41 6.40
5.19
3.92
1.73 THF-d8
OMe
O
OH
OH
O
1H NMR 300 MHz
CDCl3
12.97
8.28
1.13
0.93
f1 (ppm)
0.93
4.31
4.46
1.10
2.05
1.17
2.35
3.00
f1 (ppm)

123

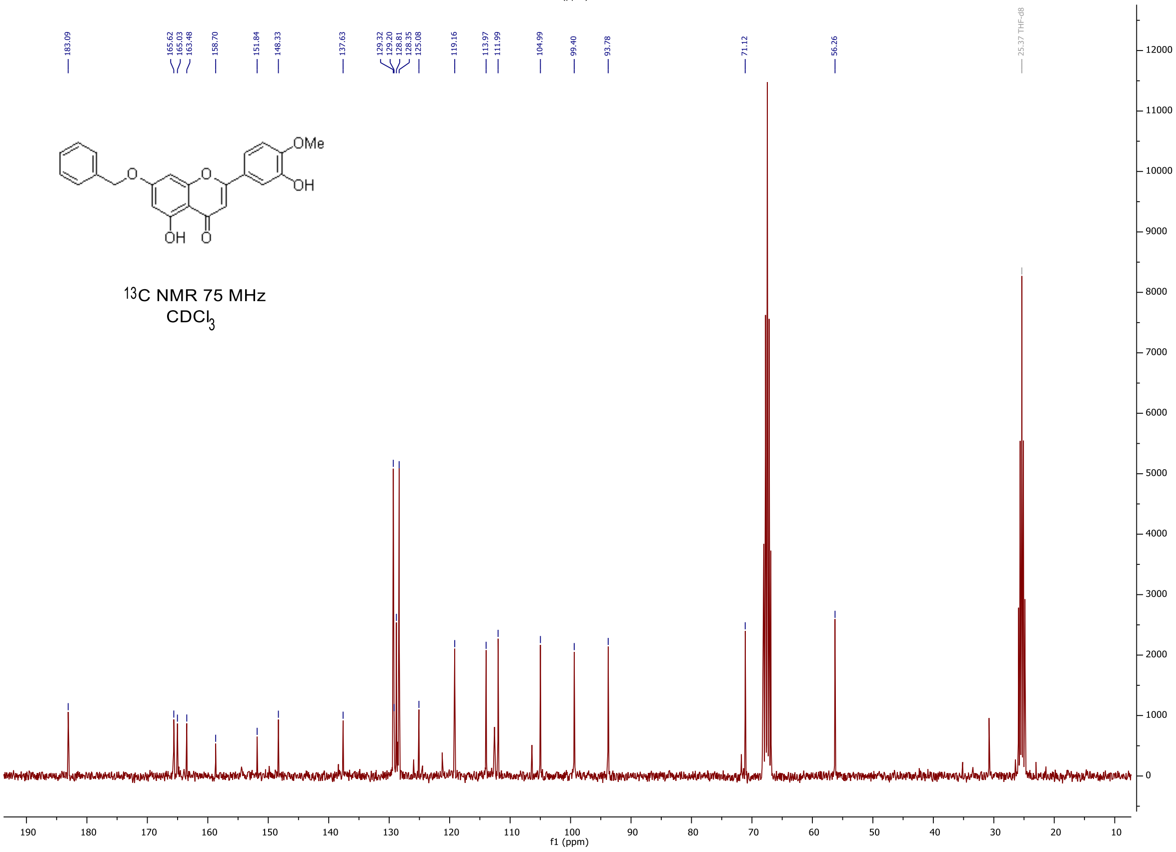
183.09
165.62 165.03 163.48
158.70
151.84
148.33
137.63
129.32 129.20 128.81 128.35 125.08
119.16
113.97
111.99
104.99
99.40
93.78
71.12
56.26
25.37 THF-d8
OMe
O
OH
OH
O
13C NMR 75 MHz
CDCl3
f1 (ppm)

124

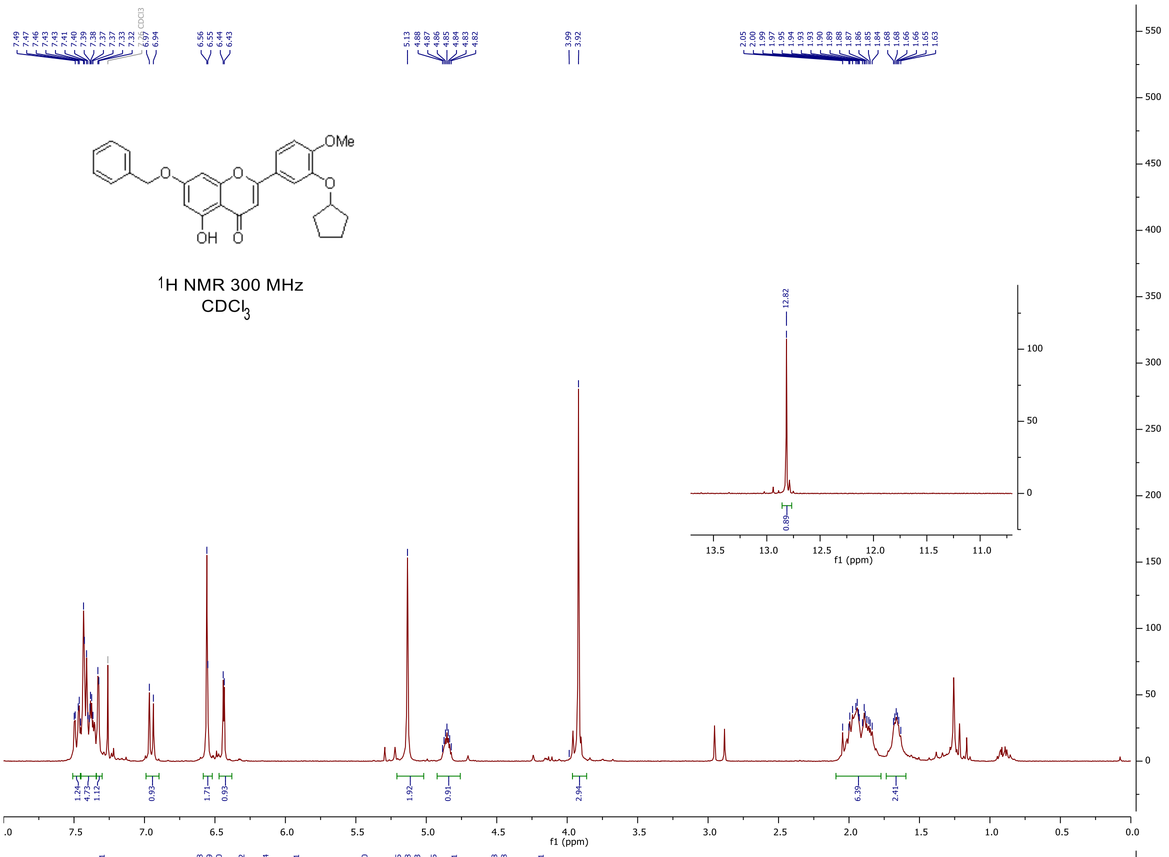


125

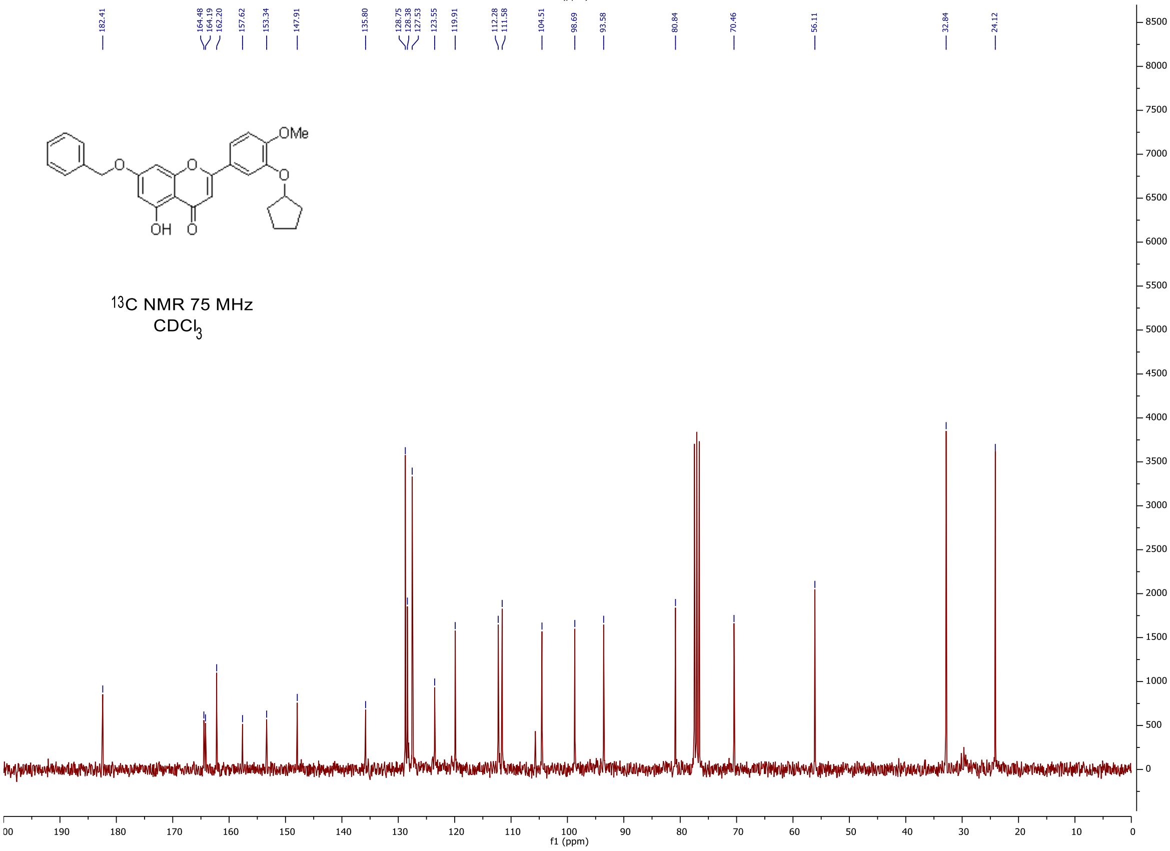


126

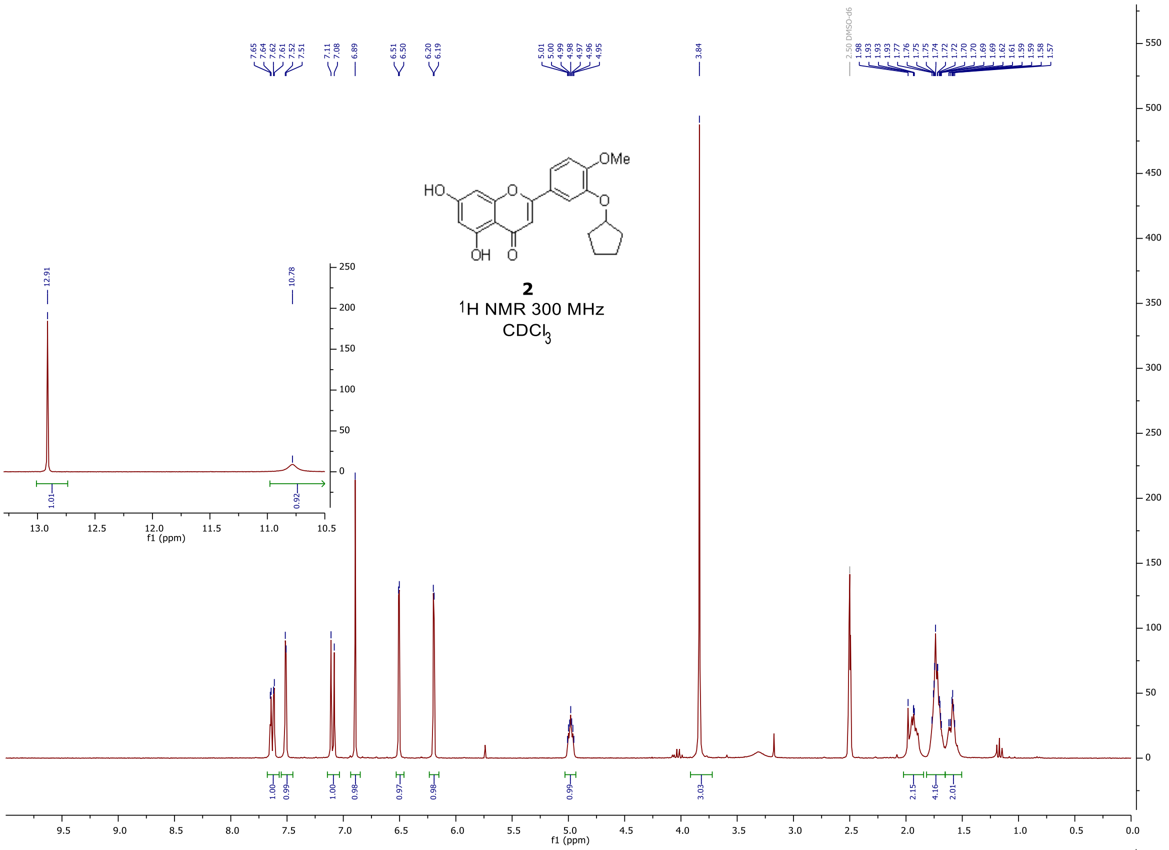
OMe
HO
O
OH
2
1H NMR 300 MHz
CDCl3
f1 (ppm)

127

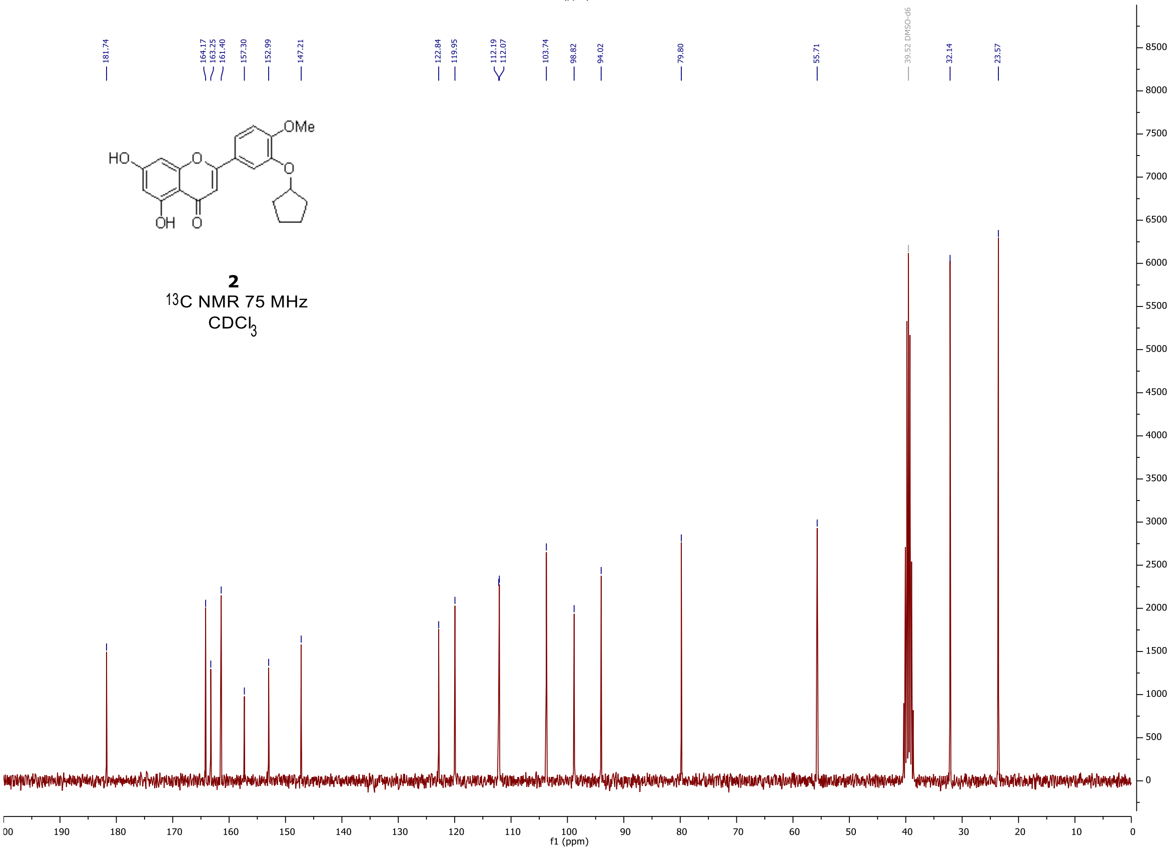
OMe
HO
O
OH
2
13C NMR 75 MHz
CDCl3
f1 (ppm)

128

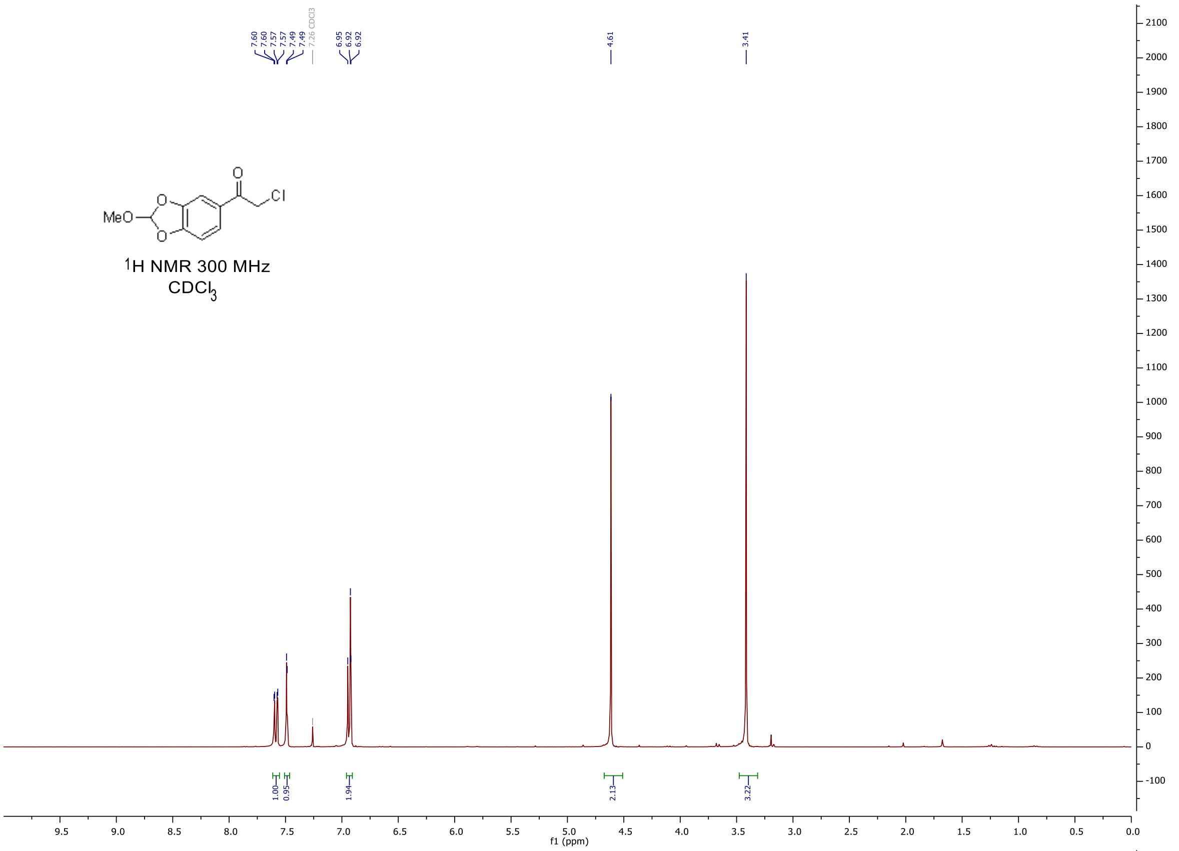

7.60
7.60
7.57
7.57
7.49
7.49
7.26 CDCl3
6.95
6.92
6.92
4.61
3.41
MeO
O
O
O
Cl
1H NMR 300 MHz
CDCl3
1.00
0.95
1.94
2.13
3.22
f1 (ppm)


129

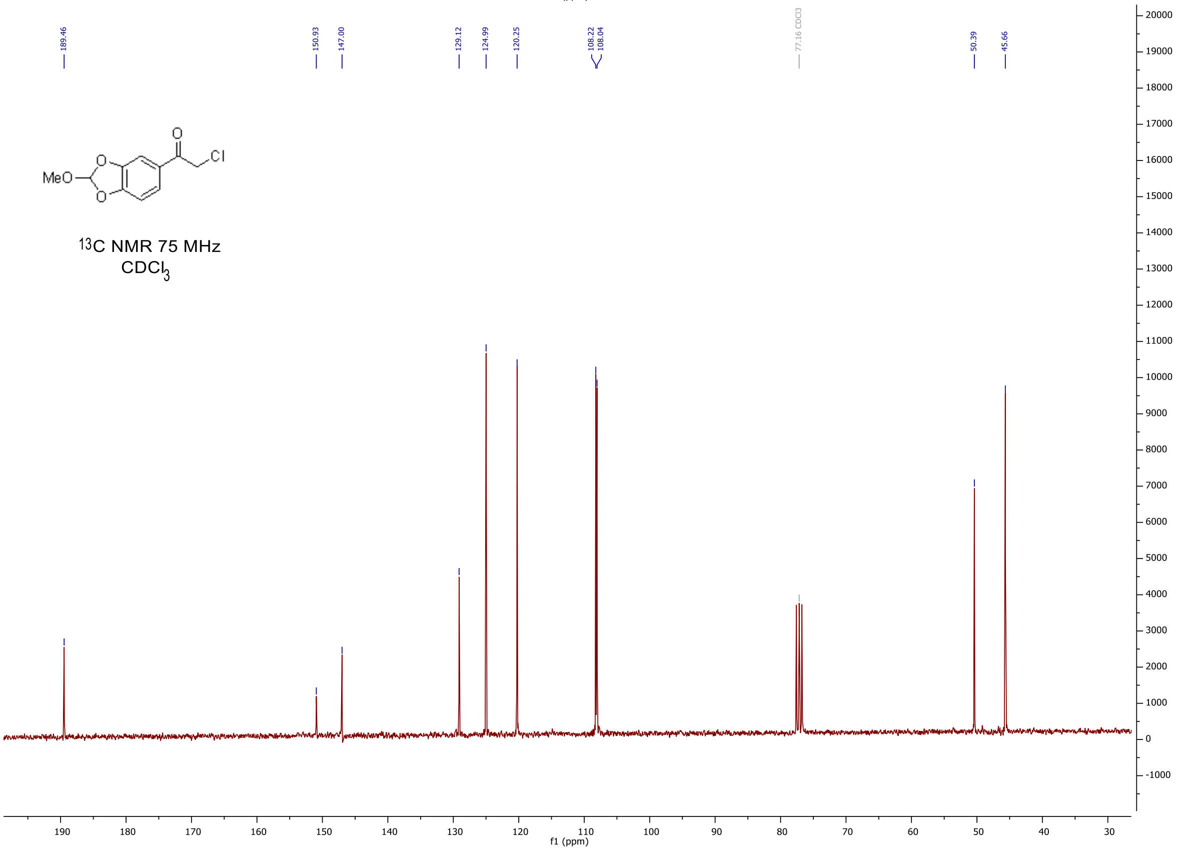

189.46
150.93
147.00
129.12
124.99
120.25
108.22
108.04
77.16 CDCl3
50.39
45.66
MeO
O
O
O
Cl
13C NMR 75 MHz
CDCl3
f1 (ppm)


130

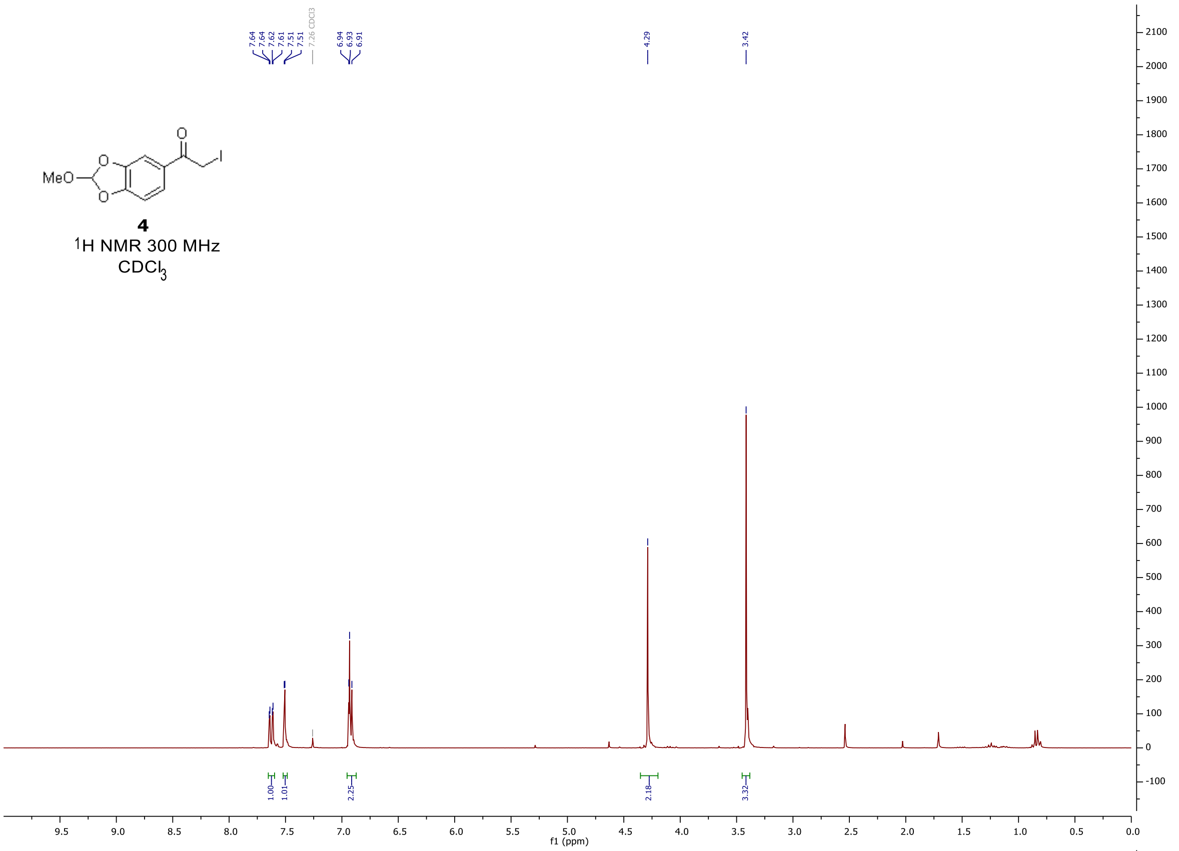
7.64
7.64
7.62
7.61
7.51
7.51
7.26 CDCl3
6.94
6.93
6.91
4.29
3.42
MeO
O
O
O
I
4
1H NMR 300 MHz
CDCl3
1.00
1.01
2.25
2.18
3.32
f1 (ppm)

131

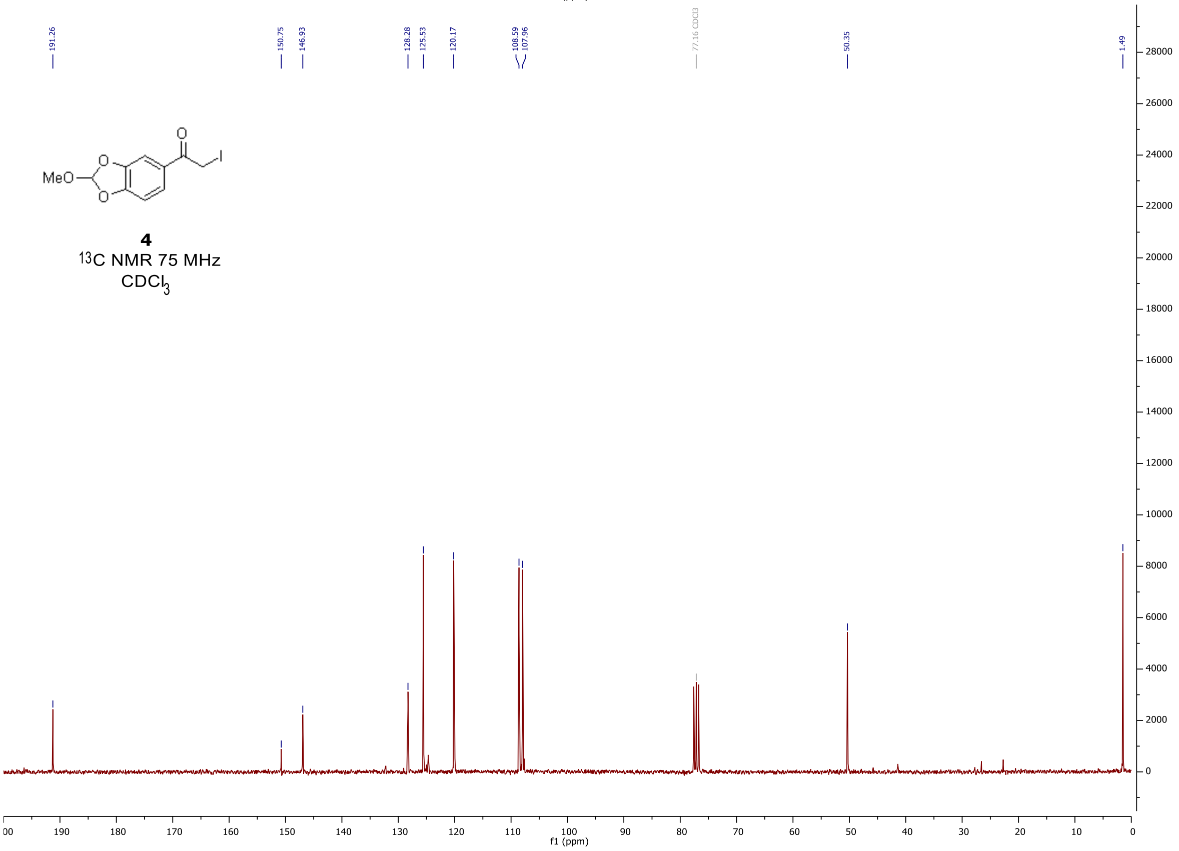
191.26
150.75
146.93
128.28
125.53
120.17
108.59
107.96
77.16 CDCl3
50.35
1.49
MeO
O
O
O
I
4
13C NMR 75 MHz
CDCl3
f1 (ppm)

132

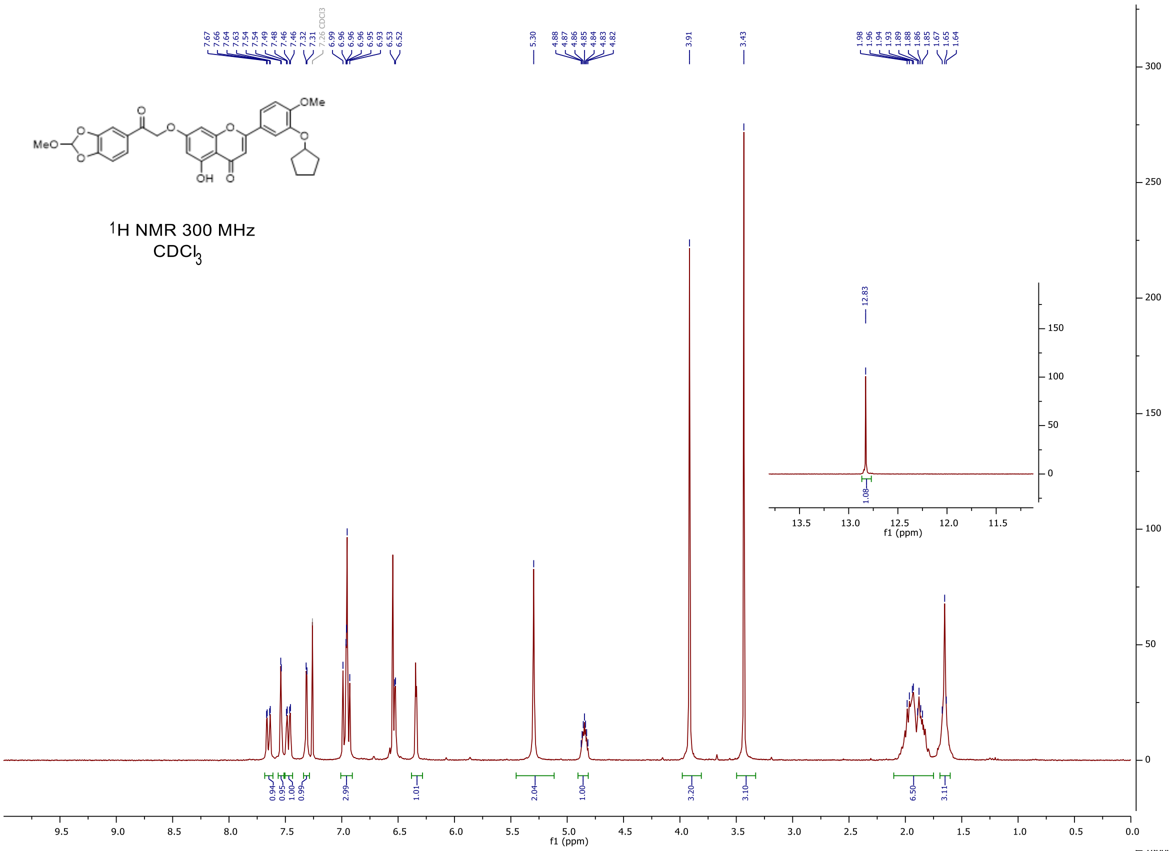


133

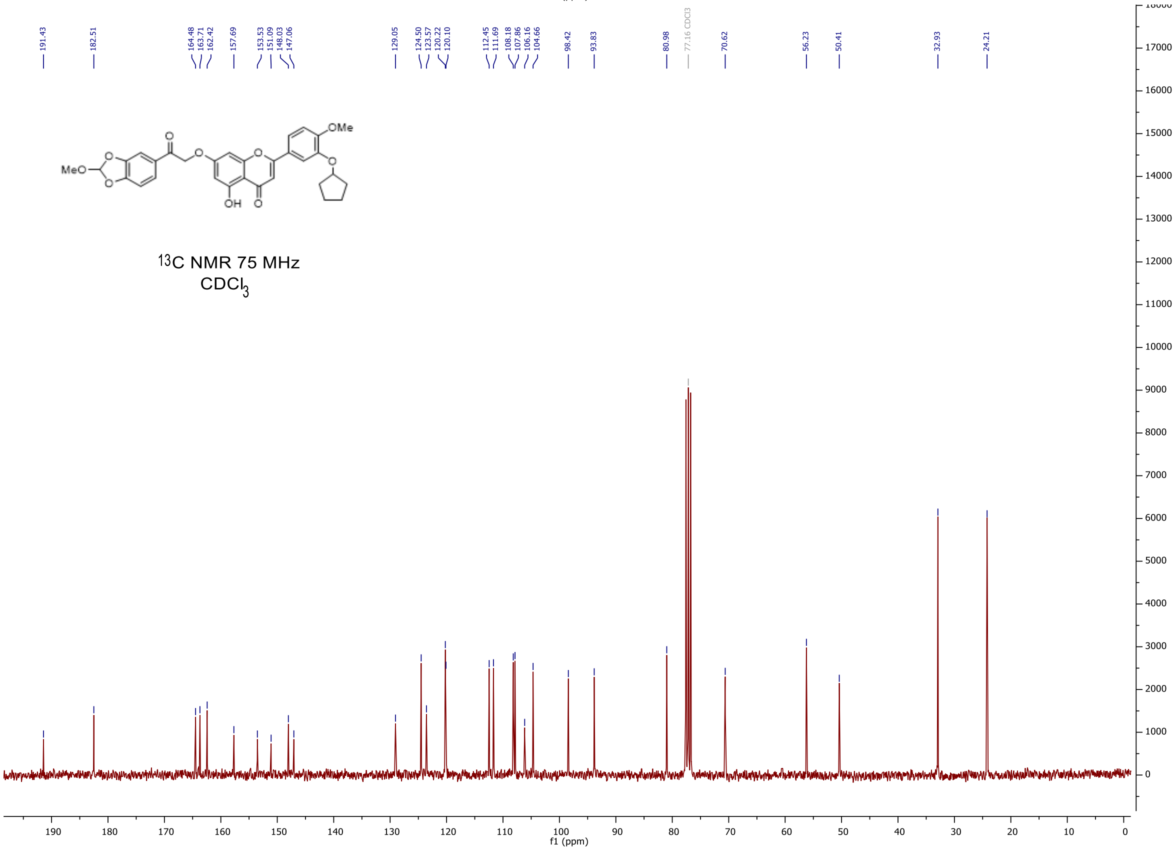


134

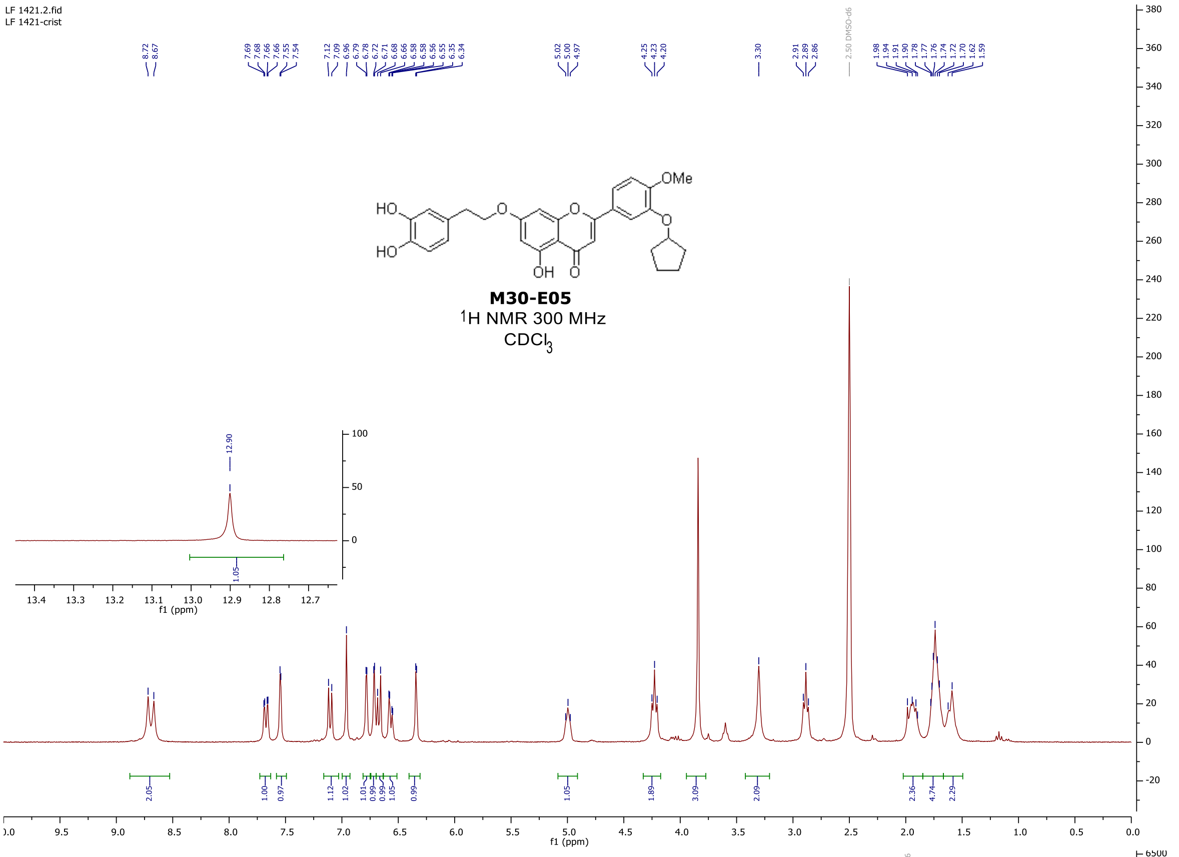
LF 1421.2.fid
LF 1421-crist
8.72 8.67
7.69 7.68 7.66 7.66 7.55 7.54
7.12 7.09 6.96 6.79 6.78 6.72 6.71 6.68 6.66 6.58 6.58 6.56 6.55 6.35 6.34
5.02 5.00 4.97
4.25 4.23 4.20
3.30
2.91 2.89 2.86
2.50 DMSO-d6
1.98 1.94 1.91 1.90 1.78 1.77 1.76 1.74 1.72 1.70 1.62 1.59
HO
HO
OMe
OH
M30-E05
1H NMR 300 MHz
CDCl3
12.90
f1 (ppm)
135

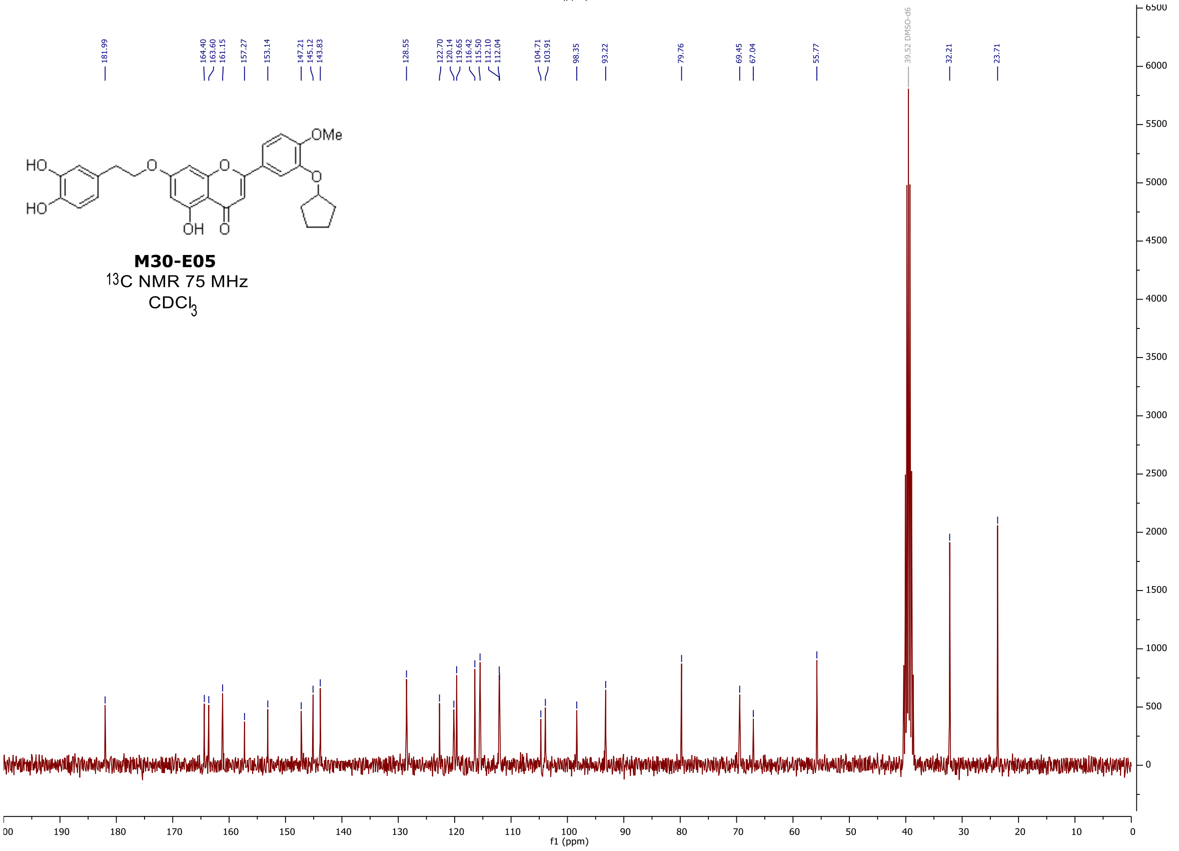
181.99
164.40 163.60 161.15
157.27
153.14
147.21 145.12 143.83
128.55
122.70 120.14 119.65 116.42 115.50 112.10 112.04
104.71 103.91
98.35
93.22
79.76
69.45 67.04
55.77
39.52 DMSO-d6
32.21
23.71
HO
HO
OMe
OH
M30-E05
13C NMR 75 MHz
CDCl3
f1 (ppm)
136
