## Supplementary figures and images for "Engineered flavonoid disrupts mitochondrial AIF/CHCHD4 complex for targeted cancer therapy"

### sup fig2

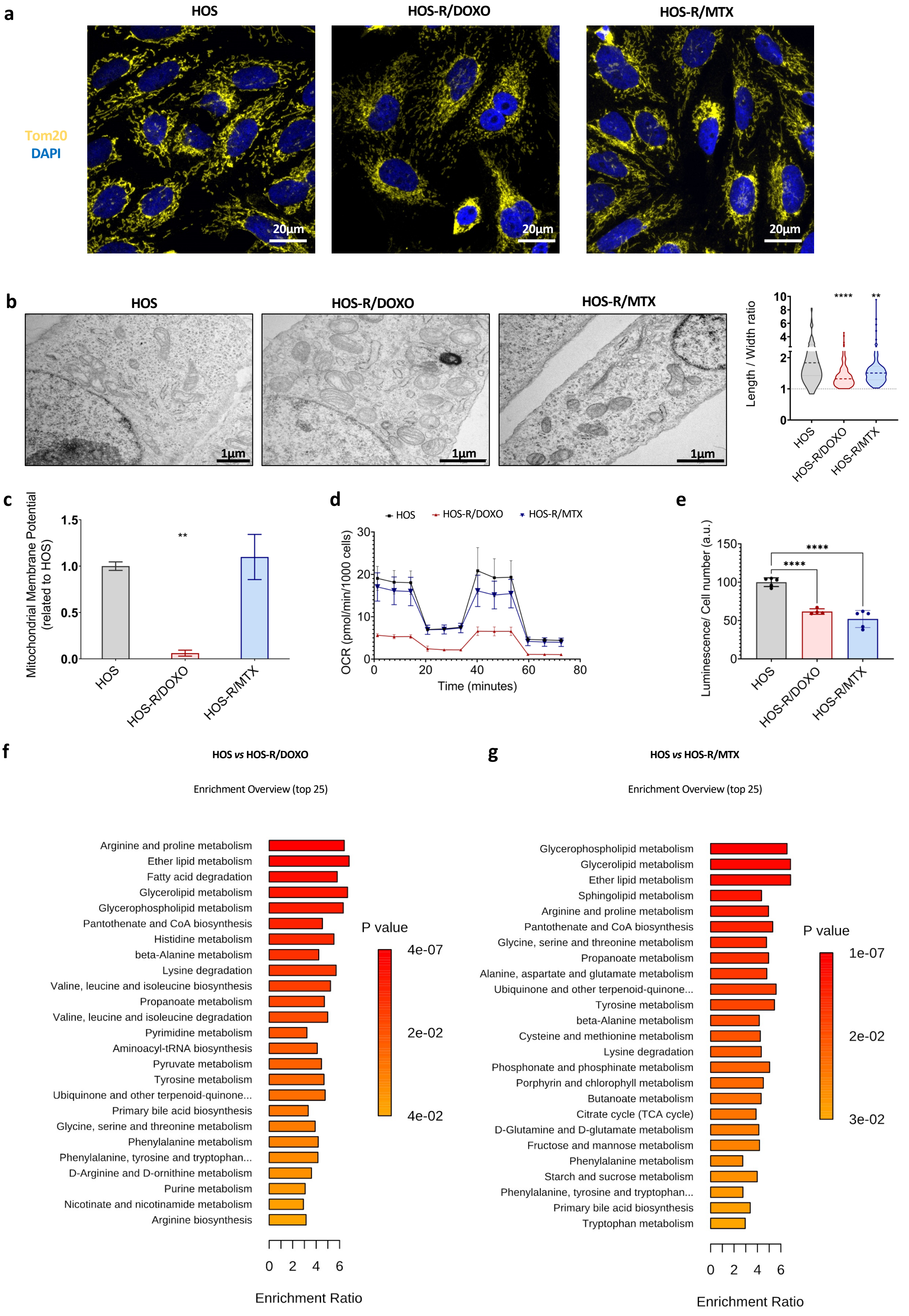


#### Supplemental Figure 2

### sup fig4

a

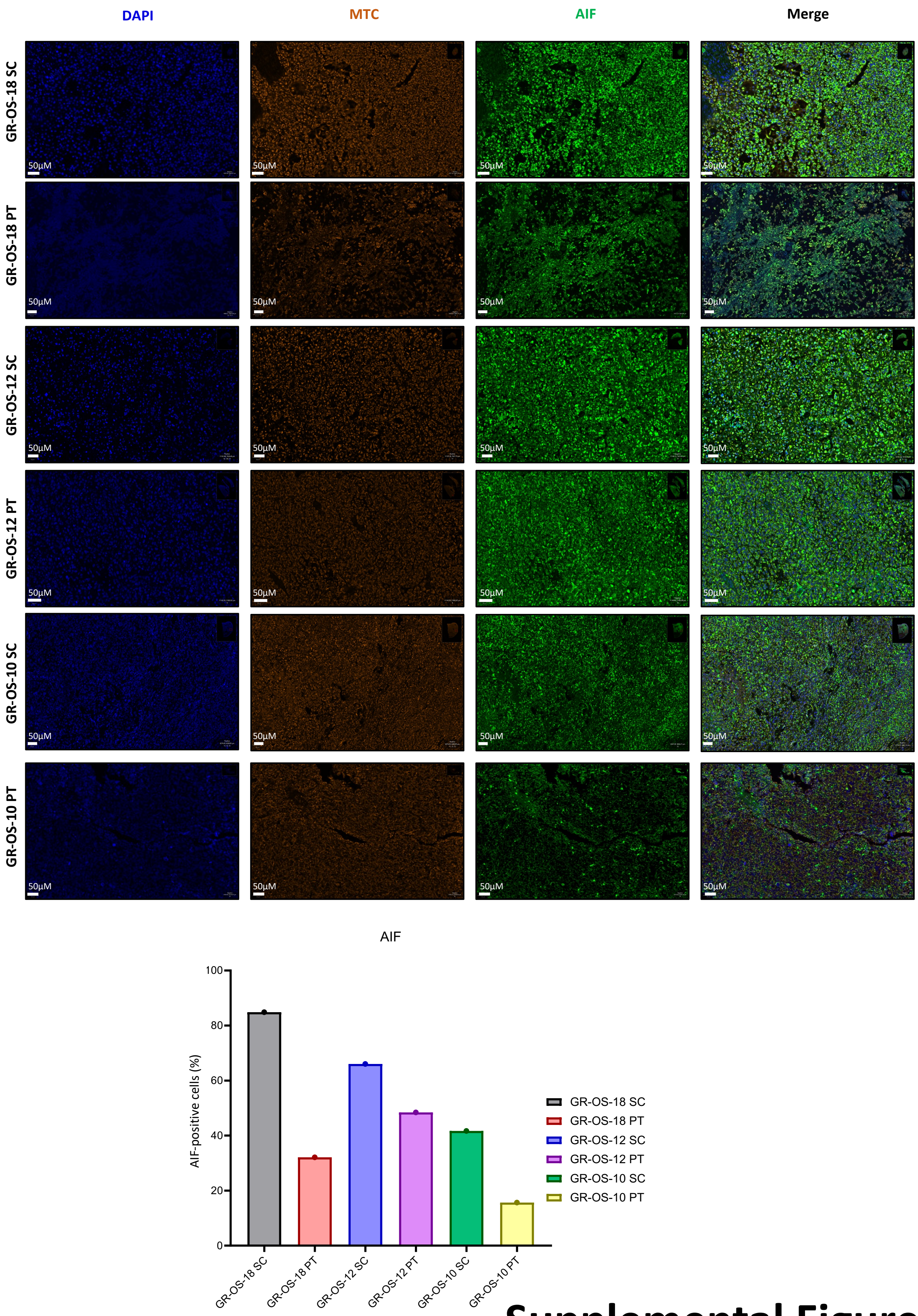


**Supplemental Figure 4a**

**b**

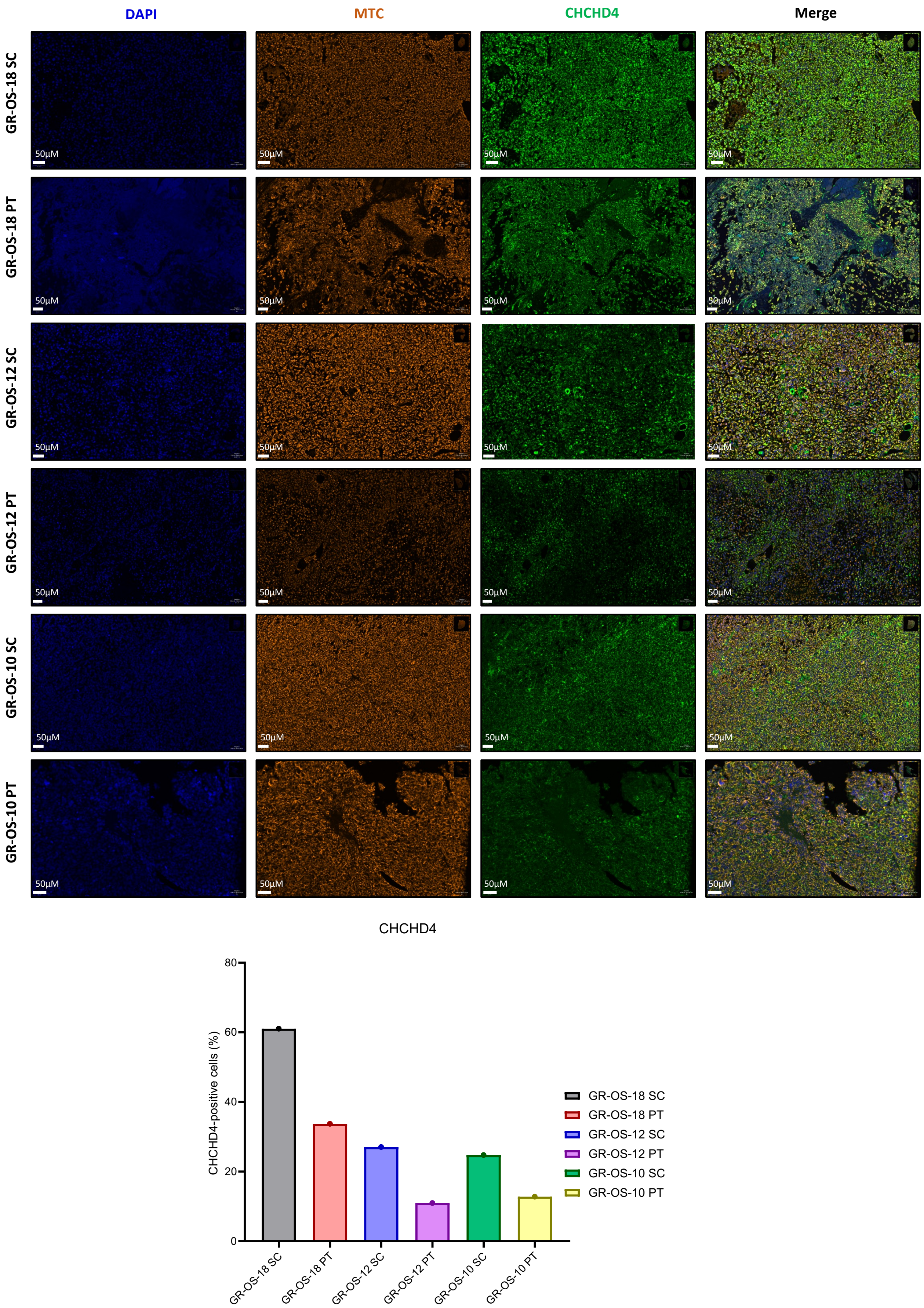


#### Supplemental Figure 4b

c

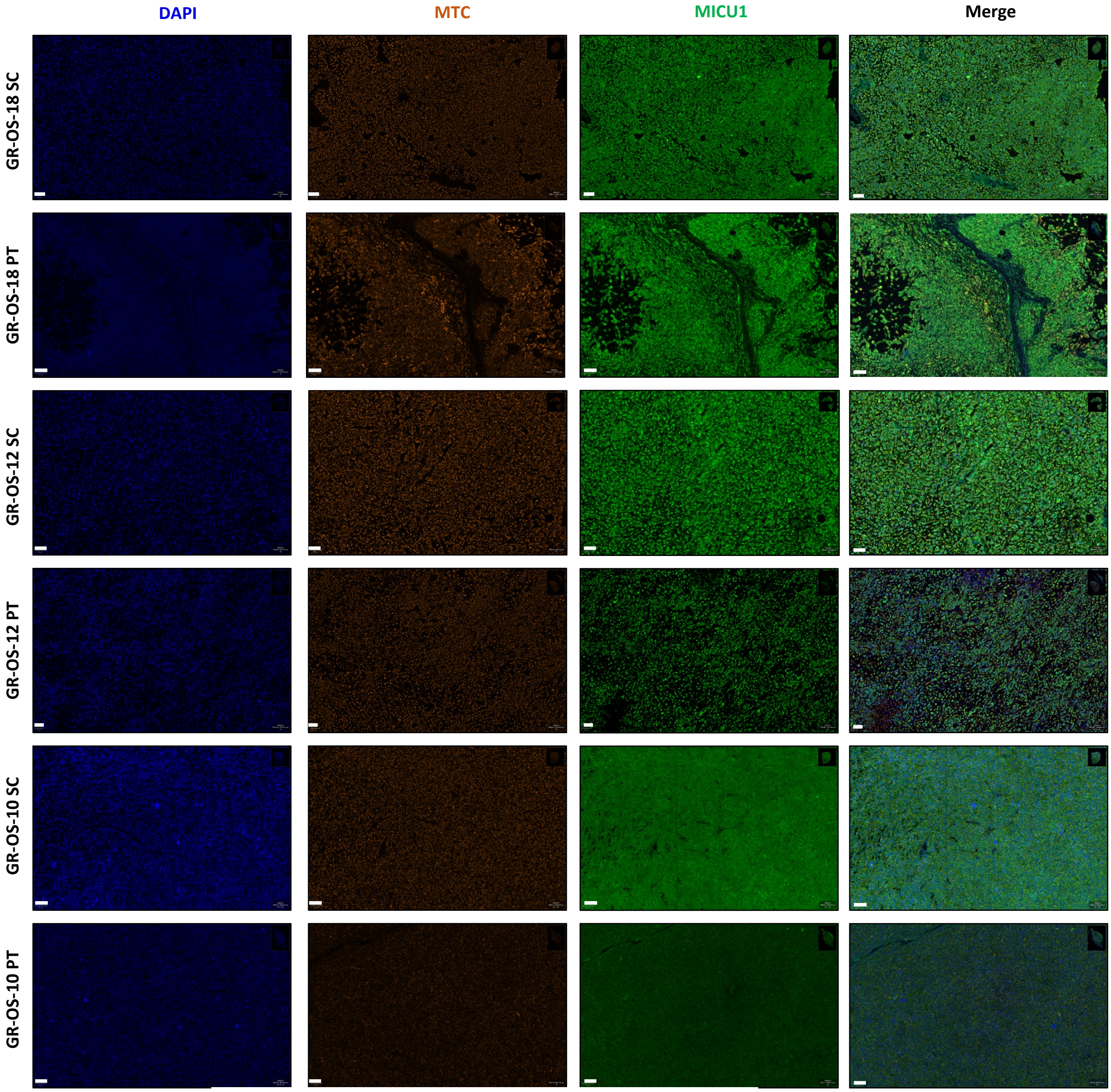


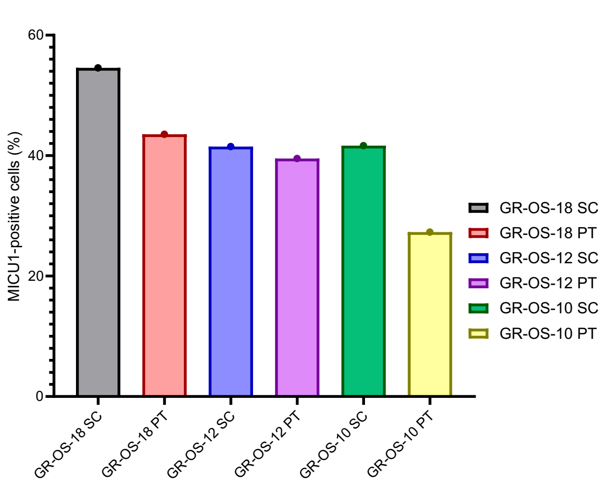


#### Supplemental Figure 4c

### supp fig1

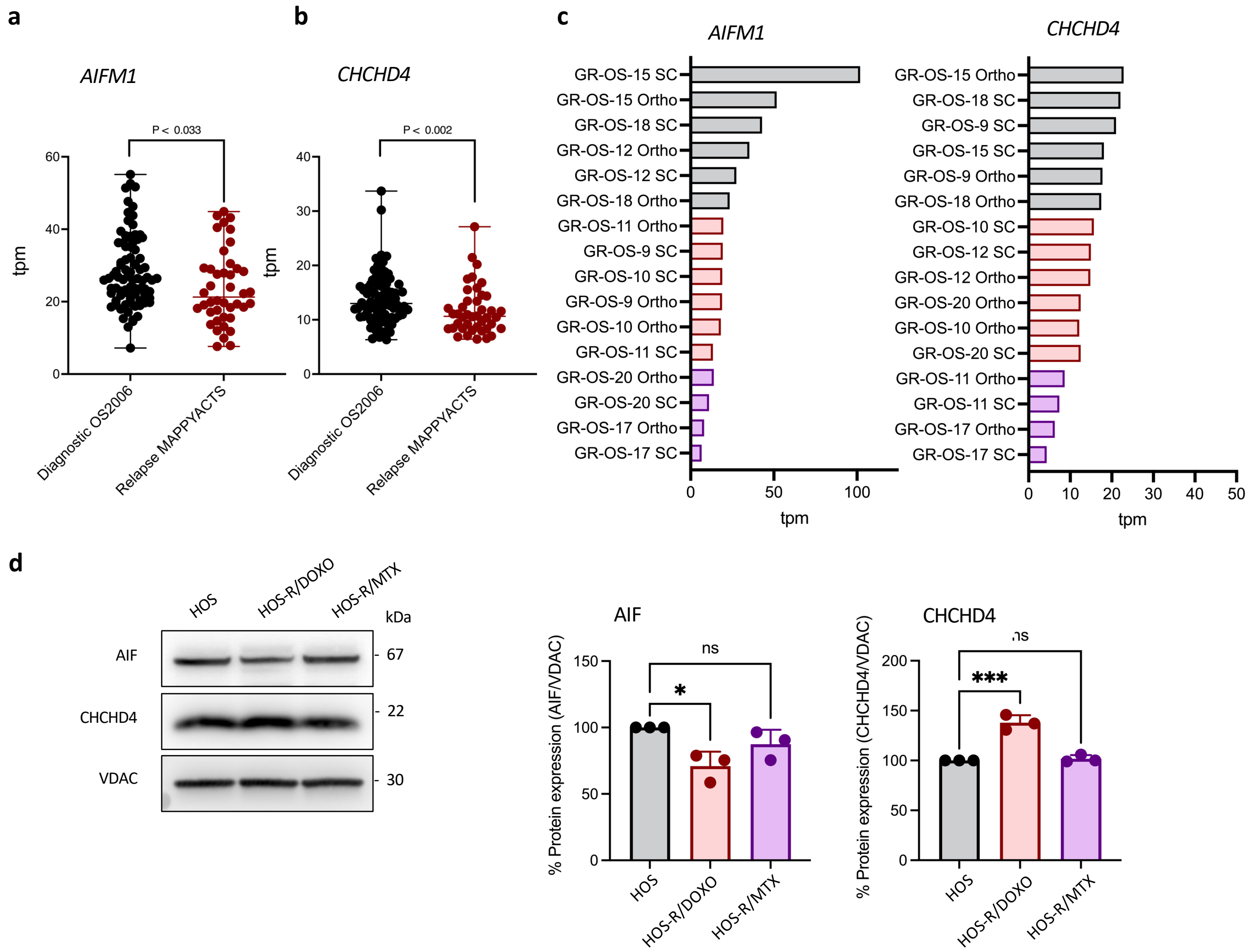


#### Supplemental Figure 1
